# Dual-targeting snRNA gene therapy rescues STMN2 and UNC13A splicing in TDP-43 proteinopathies

**DOI:** 10.64898/2025.12.01.691001

**Authors:** Trent A. Gomberg, Sara Elmsaouri, Hema M. Kopalle, Michael W. Baughn, Melinda S. Beccari, Melissa McAlonis-Downes, Jonathan W. Artates, Deepak Pant, Harriet Mak, Aaron A. Smargon, Tynan C. Sander, Esaul Garcia, Dominic P. Lee, Don W. Cleveland, Gene W. Yeo

## Abstract

Amyotrophic lateral sclerosis (ALS) is a neurodegenerative disorder caused by the selective deterioration of motor neurons in the central nervous system (CNS). A key driver of this pathogenesis is nuclear loss of ALS-associated protein TDP-43, leading to mis-splicing of TDP-43 targets including important neuronal genes *STMN2* and *UNC13A*. Here, we have developed a gene therapy strategy for ALS and related TDP-43 proteinopathies, to correct mis-splicing of both *STMN2* and *UNC13A* cryptic exons using small nuclear RNAs (snRNAs) encoded from a single vector. We identified promoter sequence elements to increase therapeutic snRNA expression by 10-fold, then further optimized the expression cassette with combinatorial snRNA targeting to rescue multiple cryptic splicing targets. The engineered snRNAs restored normal pre-mRNA processing of both *STMN2* and *UNC13A* transcripts despite TDP-43 loss of function, rescuing stathmin-2 protein levels in iPSC derived motor neurons, restoring their axonal regeneration capacity to wild-type levels. In addition, adeno-associated virus (AAV) delivery of the snRNAs to the murine central nervous system in the constitutive cryptic splicing model *Stmn2*^HumΔGU^ fully restored cortical *Stmn2* pre-mRNA processing, highlighting the utility of snRNAs as a therapeutic modality *in vivo*. Together, this study demonstrates that snRNAs are a promising and versatile therapeutic strategy for the simultaneous correction of multiple aberrant transcripts affected by cryptic splicing in TDP-43 proteinopathies.

## Introduction

Amyotrophic lateral sclerosis (ALS) is a fatal neurodegenerative disorder characterized by the loss of motor neurons in the central nervous system (CNS). This leads to rapidly progressive muscle atrophy and death typically within 2-5 years after symptom onset. Currently, ∼30,000-40,000 patients are living with ALS in the United States, and ∼5,000 individuals are newly diagnosed each year. Only ∼15% of patients present with inherited (familial) or spontaneous genetic mutations in the small set of highly penetrant ALS risk genes (*SOD1*, *C9ORF72*, *FUS*/*TLS*, and *TARDBP*/*TDP-43*[1]. For most ALS patients, no genetic cause of disease is known. Instead, the vast majority of ALS is characterized by cytoplasmic accumulation and aggregation of TDP-43, an RNA binding protein (RBP) that regulates pre-mRNA processing and metabolism, in upper and lower motor neurons [2]. This aggregation is present in >95% of post mortem ALS patient nervous system tissue [3, 4] and in other neurodegenerative disorders such as TDP-43 related frontotemporal dementia (FTD-TDP) [3, 4], Alzheimer’s disease (AD) [5–12], Parkinson’s disease (PD) [13, 14], and predominant age-related TDP-43 encephalopathy neuropathologic change (LATE-NC) [12, 15–19]. A major focus of ALS therapeutics is to correct pathology and RNA changes resulting from TDP-43 nuclear loss and subsequent aggregation [20].

An increasing body of evidence demonstrates an important role of TDP-43 within neurons is to suppress incorporation of so-called “cryptic exons” into nascent mRNAs [21–29]. These cryptic mis-splicing events occur upon nuclear depletion of TDP-43 and appear to be major contributors to disease pathogenesis and dysregulation of RNA metabolism. Stathmin-2 (*STMN2*) and *UNC13A* mis-splicing have emerged as prominent neuronal pre-mRNAs normally guarded from cryptic exon incorporation by TDP-43 binding, and are promising therapeutic targets for targeted RNA restoration.

*STMN2* is among the top twenty most highly expressed transcripts in motor neurons [28] and is essential for axon integrity, neuromuscular junction maintenance, and regenerative capacity [28–32]. It is thought that the *STMN2* mRNA is the most sensitive to TDP-43 depletion [12, 28]. Knock-down of TDP-43 leads to a commensurate or greater depletion of *STMN2* mRNA relative to *TARDBP* mRNA [12, 28, 33]. This is regulated through several tandem UG hexamers comprising 24 nucleotides in total between exons 1 and 2 of *STMN2* pre-mRNA which comprise a TDP-43 binding site [28, 33]. In healthy neurons, TDP-43 binds and suppresses recognition of the 3’ splice site of a cryptic exon (exon 2a; Fig. 1a), whose incorporation and premature poly-adenylation truncates the *STMN2* mRNA transcript, depleting functional stathmin-2 (Fig. 1b).

**Figure 1:**
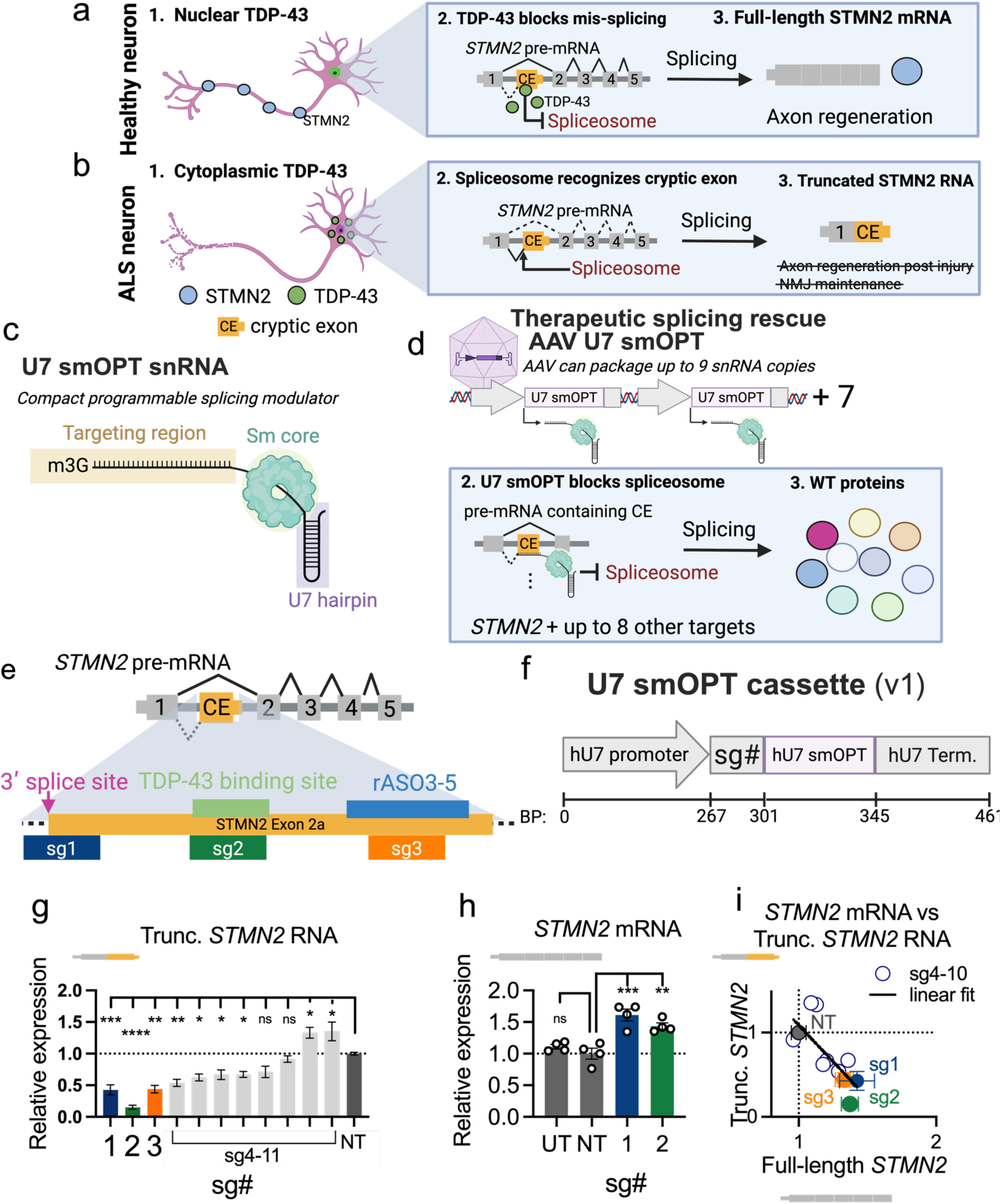
Targeted U7smOPT suppresses cryptic splicing in ALS model. a) Diagram depicting TDP-43’s function in splicing in healthy motor neurons. TDP-43 binds to intronic regions in *STMN2* pre-mRNA and blocks spliceosome recruitment, preventing inclusion of the cryptic exon (CE, exon 2a). b) Diagram showing the events in TDP-43 proteinopathies, where the protein aggregates in the cytoplasm. TDP-43’s nuclear depletion leads to the inclusion of the *STMN2* cryptic exon, and depletion of full-length *STMN2* mRNA, which is essential for axon regeneration and neuromuscular junction (NMJ) maintenance. c) Schematic depicting U7smOPT, a small nuclear RNA with an antisense region for pre-mRNA targeting, an Sm core binding motif (SmOPT), and a U7 snRNA hairpin. d) Diagram illustrating a potential multi-valent gene therapy vector which contains up to nine U7smOPT expression cassettes that suppress multiple cryptic splicing events. e) Schematic showing the targeting strategy employed to block inclusion of the *STMN2* cryptic exon. The 3’ splice site, TDP-43 binding site, and the targeting site of rASO3-5 in Baughn et al 2023 are shown. Guides of 25 or 36 base pairs were tiled along these regions. Guides 1-3 (sg1, sg2, sg3) are depicted f) Schematic of the DNA expression cassette of U7smOPT with the human U7 (hU7) promoter and terminator (Term.), with a *STMN2* targeting guide (sg#) and a human U7smOPT backbone. g,h) Bar plots of levels of truncated *STMN2* (g) or full-length *STMN2* (h) mRNA in SH-SY5Y^N352S/N352S^ cells transduced with the indicated snRNAs targeting the *STMN2* cryptic exon region or with a non-targeting snRNA (NT), or from untreated cells (UT). sg1-3 are in color (blue, green and orange, respectively). Data are rt-qPCR of target RNA normalized to GAPDH (ΔC_q_), relative to non-targeting snRNA (ΔΔC_q_). Statistical tests are one-way ANOVA with Dunnett’s correction. ****P < 0.0001; ***P < 0.001; **P < 0.01; *P < 0.05; ns, not significant. (g) N=2 biological replicates. (h) N=4 biological replicates. i) Scatter plot comparing levels of truncated *STMN2* mRNA levels (y-axis) to full-length *STMN2* mRNA (x-axis). Error bars are only included for NT and sg1-3. R^2^ = 0.51. N=2 biological replicates.

*UNC13A* is a highly conserved protein that is critical for synaptic vesicle release. Single nucleotide variants in *UNC13A* are tightly associated with ALS in GWAS studies [34, 35]. Like intron 1 of *STMN2*, *UNC13A* contains a stretch of GU-rich TDP-43 binding motif sequences between exons 20 and 21. In the absence of TDP-43 one of two cryptic isoforms are produced, but both lead to nonsense mediated decay of the transcript, depleting UNC13A protein levels [24, 25].

The current state of the art for targeted therapeutic splicing correction is splice switching antisense oligonucleotides (SSOs), including the founding example that is now a standard of care for the fatal childhood motor neuron disease spinal muscular atrophy (SMA) [36–38]. Several academic and commercial groups have developed splice switching antisense oligonucleotides (SSO) to rescue endogenous processing of *STMN2* and *UNC13A* [33, 39], and both approaches are currently in clinical trials. While SSOs have shown spectacular clinical success in SMA, they require repeat dosing, and multi-target combinational ASO therapy remains untested.

To develop a single-dose multivalent pre-mRNA corrective therapy targeting RNA metabolism defects of TDP-43 proteinopathy, the modified RNA-guided U7smOPT is a promising option [40]. This reprogrammed small RNA is derived from the U7 small nuclear RNA (snRNA) involved in histone tail processing. U7 canonically binds the LSM protein ring and to the tail of histone mRNA. In U7smOPT, the LSM binding site is replaced with an Sm core binding site [41] (Fig. 1c). With this modification, U7smOPT is more stable than U7 snRNA and no longer participates in histone tail processing [42]. Typically, U7smOPT has been used to target disease causing mutations to correct splicing at those loci [43–45], though their applications have expanded from RBP blocking to exon inclusion, A to I editing and pseudouridylation [46, 47].

Unlike ASOs, U7smOPT is encodable in lentiviral or AAV vectors. This combines the advantages of competing pre-mRNA targeting technologies, ASOs and RNA-targeting CRISPR effectors. First, up to nine distinct U7smOPTs can be packaged in a single adeno-associated virus (AAV) capsid [48] for durable long-term expression (Fig. 1d). Second, the U7smOPT relies only upon endogenous RBPs, requiring no foreign protein delivery that could elicit an immune response [42]. Third, similar to the case of ASOs for treatment of SMA, U7smOPT has been used in patients with promising preliminary data showing restoration of full-length dystrophin [49, 50]. These attributes make U7smOPT an attractive therapeutic option for correcting mis-splicing in TDP-43 proteinopathies like ALS.

In this study, we develop an snRNA-based gene therapy approach for long-term multi-valent suppression of TDP-43 dependent cryptic splicing events of the human *STMN2* and *UNC13A* genes.

## Results

### U7smOPT snRNAs block cryptic splicing in *STMN2* and *UNC13A*

We identified three promising regions within the *STMN2* cryptic exon to target with U7smOPT based on three different strategies for blocking splicing. From 5’ to 3’, the first region we identified was the 3’ splice acceptor site (3’ SS), which is essential for cryptic exon recognition and has been successfully leveraged for suppressing splicing in the literature (magenta; Fig 1e)[33, 51]. Second, we took inspiration from TDP-43’s endogenous function, to block splicing through steric hinderance, and targeted the TDP-43 binding sites (green; Fig 1e)[33]. Lastly, we designed guides to target the same region identified in Baughn et al, where 250 ASOs were screened for cryptic splicing suppression. Three of the five lead ASOs identified in that study targeted between 99-126 base pairs downstream of the 3’ SS of the cryptic exon (blue, Fig. 1e).

We tested 10 U7smOPT antisense guide sequences (guides) by delivering lentivirus encoding snRNA expressed off the human U7 promoter and terminator to an SH-SY5Y neuroblastoma cell line homozygous for the TDP-43^N352S/N352S^ mutation (Fig. 1f; SH-SY5Y^N352S/N352S^; [28]). These cells express approximately 50% of the full-length STMN2 mRNA and 10x the cryptically spliced truncated *STMN2* RNA as wild-type SH-SY5Y cells.

Of the ten guides tested, three significantly suppressed truncated *STMN2* snRNA in an initial screen (Fig. 1g; p < 0.05; 15%-45% expression relative to NT snRNA; N=2). For the two best performing snRNAs, sg1 and sg2, full-length *STMN2* mRNA increased as well (Fig. 1h; 144%-sg1; sg2 - 129). Altogether, full-length *STMN2* mRNA increased linearly with a decrease in truncated *STMN2* RNA, confirming that these snRNAs are reducing the truncated transcript and restoring the full-length transcript through splicing suppression (Fig. 1i; R^2^=0.54; linear fit: Y intercept = 2.93, Slope = -1.864). We moved forward with sg1 as our lead candidate, though we continued with sg2 as well for some experiments.

### Optimized snRNA expression cassette produces an order of magnitude more snRNA

With effective U7smOPT snRNAs for *STMN2* cryptic exon suppression, we next sought to optimize these snRNAs for a downstream application in adeno-associated virus (AAV) gene therapy. We wanted to develop a vector that expressed snRNAs efficiently and would allow us to deliver several snRNAs with a single vector. The latter can be limited by the inherent nature of sequences with a high percentage of homology to recombine and lead to inefficiencies in AAV packaging. Thus, we optimized snRNA promoter-terminators (cassettes) for two parameters: sg1 expression efficiency (snRNA copies per template DNA copy) and cassette homology (sequence similarity with different snRNA cassettes) (Fig. 2a).

**Figure 2:**
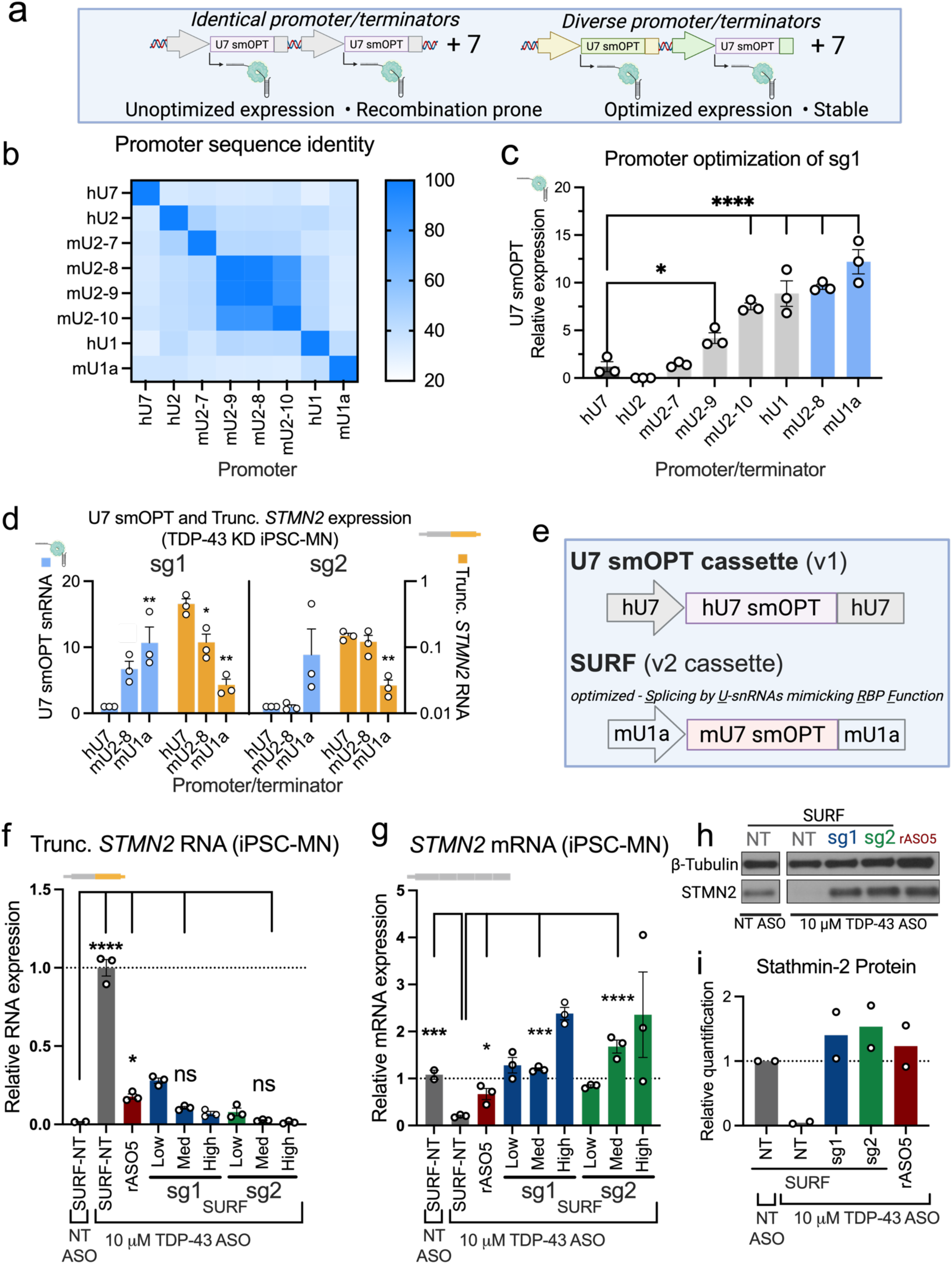
Optimized snRNA cassette corrects STMN2 splicing in mature iPSC derived motor neurons. a) Schematic representation of a hypothetical multivalent vector. b) Heatmap showing percent nucleotide identity of the promoters used in this study. c) Bar plot comparing the levels of sg1 snRNA in HEK293T cells transfected with plasmid constructs driving U7smOPT expression from the indicated promoters. Data are rt-qPCR normalized to PuroR transcript levels (ΔC_q_), relative to normalized levels of sg1 U7smOPT driven by the hU7 promoter (ΔΔC_q_). N=3 biological replicates. One-way ANOVA with Dunnett’s correction. ****P < 0.0001; *P < 0.05; not significant if not otherwise indicated. d) Bar plot comparing the levels of sg1 and sg2 U7smOPT (blue, left y-axis) or truncated *STMN2* mRNA (orange, right y-axis) in iPSC-MN transduced with lentiviral vectors driving U7smOPT expression from the indicated promoters and terminators. Data are rt-qPCR normalized to PuroR mRNA levels (ΔC_q_), relative to normalized levels of a non-targeting U7smOPT driven by the hU7 promoter (ΔΔC_q_). N=3 biological replicates. One-way ANOVA with Dunnett’s correction. hU7 was used as a control in each comparison. **P < 0.01; *P < 0.05; not significant unless otherwise indicated. e) Schematic representation of the v1 U7smOPT cassette driven by the human U7 promoter/terminator (top) and the optimized design we term SURF (optimized Splicing by U-snRNA mimicking RBP function; bottom), which encompasses sg1 or sg2 expressed under an mU1a cassette. f,g) Bar plots showing levels of truncated *STMN2* (f) or full-length *STMN2* (g) mRNA in day-42 iPSC-MN transduced with lentiviral vectors expressing the indicated SURF constructs from an mU1a promoter/terminator at the indicated relative titers (low = 1×, mid=5×, high=15×; non-targeting (NT) SURF construct titers were 5×), or treated with the *STMN2*-targeting SSO rASO5. Cells were pretreated with either a non-targeting (NT) ASO or a TDP-43 targeting ASO, as indicated. Data are rt-qPCR of the target mRNA normalized to GAPDH mRNA levels, relative to normalized levels of target mRNA in cells transduced with the SURF-NT construct and treated with either TDP-43 ASO (f) or NT ASO (g). N=2 biological replicates; SURF-NT; N=3 biological replicates; all other samples. One-way ANOVA with Dunnett’s correction. ****P < 0.0001; ***P < 0.001; *P < 0.05; ns, not significant h,i) Western blot showing levels of STMN2 in iPSC-MN transduced with lentiviral vectors expressing the indicated SURF constructs or treated with the *STMN2*-targeting SSO rASO5. Cells were pretreated with either a non-targeting (NT) ASO or a TDP-43 targeting ASO, as indicated. (i) Bar plot represents quantification of N=2 biological replicates.

Most studies that utilize U7smOPT use a human or mouse U7 cassette, though some use U1 snRNA cassettes with variable success relative to U7 [46, 47, 52, 53]. The inconsistency between different studies suggests that snRNA cassette performance is somewhat dependent on the sequence identity of the snRNA they are expressing. In addition to the standard complement of U1 and U7 cassettes, we tested human and mouse U2 promoters, utilizing the canonical U2 snRNA promoter in humans. From mice, four U2 snRNA cassettes were tested that had been identified as functional in the literature [54].

In all, we screened 7 additional cassettes from U1 and U2 genes from both mouse and human including (1) the two canonical U1 snRNA promoters from human and mouse (hU1 and mU1a), (2) the canonical human U2 promoter, and (3) four additional mouse U2 promoters identified in Jia et al 2012 [54]. Additionally, U7smOPT snRNAs expressed under mouse cassettes were modified to match the mouse U7 snRNA in sequence.

Importantly, the promoters and terminators we tested are distinct from one another, with no two promoters or terminators (outside of the mouse U2 promoters) having greater than 55% sequence homology (Fig. 2b; Fig. s3a). While the overall homology between the cassettes is low, the functional elements within the promoter have higher homology than the surrounding areas (DSE, PSE, OCTA binding site; Fig. s3b; [52, 55, 56]).

Sg1 expression from each of the seven promoter cassettes was assessed in HEK293T cells by transient transfection. The synthetic snRNA expression was assessed by anchor addition and qPCR, a common method for quantifying small RNAs (Fig. s3c). Five of the seven promoters expressed significantly higher levels of sg1 U7smOPT than the human U7 promoter-terminator, including the mU1a, hU1, and three of the four mouse U2 promoters (mU2-8, mU2-9, mU2-10)(Fig. 2c).

We then tested the two lead cassettes, mU1a and mU2-8, in a disease relevant context. We delivered sg1 and sg2 by lenti to iPSC-MN with TDP-43 knock-down. We quantified truncated *STMN2* RNA and U7smOPT by rt-qPCR. Again, we saw a significant increase in expression of sg1 under the mU1a and mU2-8 cassette compared to the hU7 cassette (sg1 expression in iPSC-MN: hU7 (1) < mU2-8 (6.68) < mU1a (10.67); Fig 2d). Increased expression of the snRNA led to greater suppression of the truncated *STMN2* RNA, as expected (trunc. *STMN2* expression in iPSC-MN: hU7 (0.45) < mU2-8 (0.11) < mU1a (0.026); Fig 2d). The same study for sg2 saw similar results for hU7 and mU1a, but the mU2-8 cassette did not express more sg2 nor suppress the truncated STMN2 RNA more than sg2 under the hU7 cassette (hU7: 0.14, mU2-8: 0.11, mU1a: 0.027; Fig. 2d, yellow). This confirmed our previously stated hypothesis that snRNA expression is somewhat dependent on sequence-cassette combinations. This suggests that for an optimal gene therapy cargo, lead candidate snRNAs should be screened with several cassettes. Lastly, we performed rolling circle amplification (RCA) of the 36 base pair guides to quantify snRNA levels. These RCA results confirmed the results observed in Figure 2 (Fig. S4). Moving forward, we will refer to the lead vector tested here, mU1a promoter driven **sg1 or sg2** with mouse U7smOPT, optimized - <u>S</u>plicing by <u>U</u>-snRNAs mimicking <u>R</u>BP <u>F</u>unction, or **SURF**. Expression cassettes are named either SURF-sg1, SURF-sg2, or SURF-NT (Fig. 2e).

### Restoration of STMN2 by SURF-sg1 and SURF-sg2

Next, we sought to restore *STMN2* mRNA and protein in a disease relevant context using our novel SURF system. To do this, we again knocked-down TDP-43 in iPSC-MN using a TDP-43 (*TARDBP*) targeting ASO (D30-42). We delivered SURF-sg1 and SURF-sg2 lentivirus to TDP-43 depleted iPSC-MN post-maturity (D32). In cells with TDP-43 (*TARDBP*) mRNA reduced almost completely, both SURF-sg1 and SURF-sg2 restored *STMN2* mRNA to wild-type or above wild-type levels while reducing cryptic exon levels more than 90% (Fig. 2f-g; Fig. s5a). Interestingly, SURF-sg2 seemed to outperform SURF-sg1 in cryptic exon suppression, and so we compared multiple does of each construct. We found that SURF-sg2 more efficiently suppresses the cryptic exon, but that there is no significant difference between the restoration of *STMN2* mRNA between the two constructs at multiple doses (Fig. s5b-c). Additionally, STMN2 protein was restored to wild-type or above wild-type levels, even outperforming the state-of-the-art SSO, rASO5 (Fig. 2i).

### Rescue of axon regeneration by SURF-sg1

The STMN2 protein is essential for axon plasticity and regeneration. While the molecular pathology of ALS is difficult to recapitulate in vitro, one of the more striking pathologies observed in TDP-43 (and so STMN2 protein) depleted iPSC-MN is the stunted regrowth of motor axons that have been severed [28, 30]. To demonstrate functional STMN2 protein restoration, we performed an axotomy and regrowth assay in iPSC-MN with TDP-43 knock-down treated with either SURF-sg1 or SURF-NT (Fig. 3a). After mechanical axotomy at day 42, cells were cultured until day 56 where the sustained restoration of STMN2 protein was confirmed by IF (Fig. 3b).

**Figure 3:**
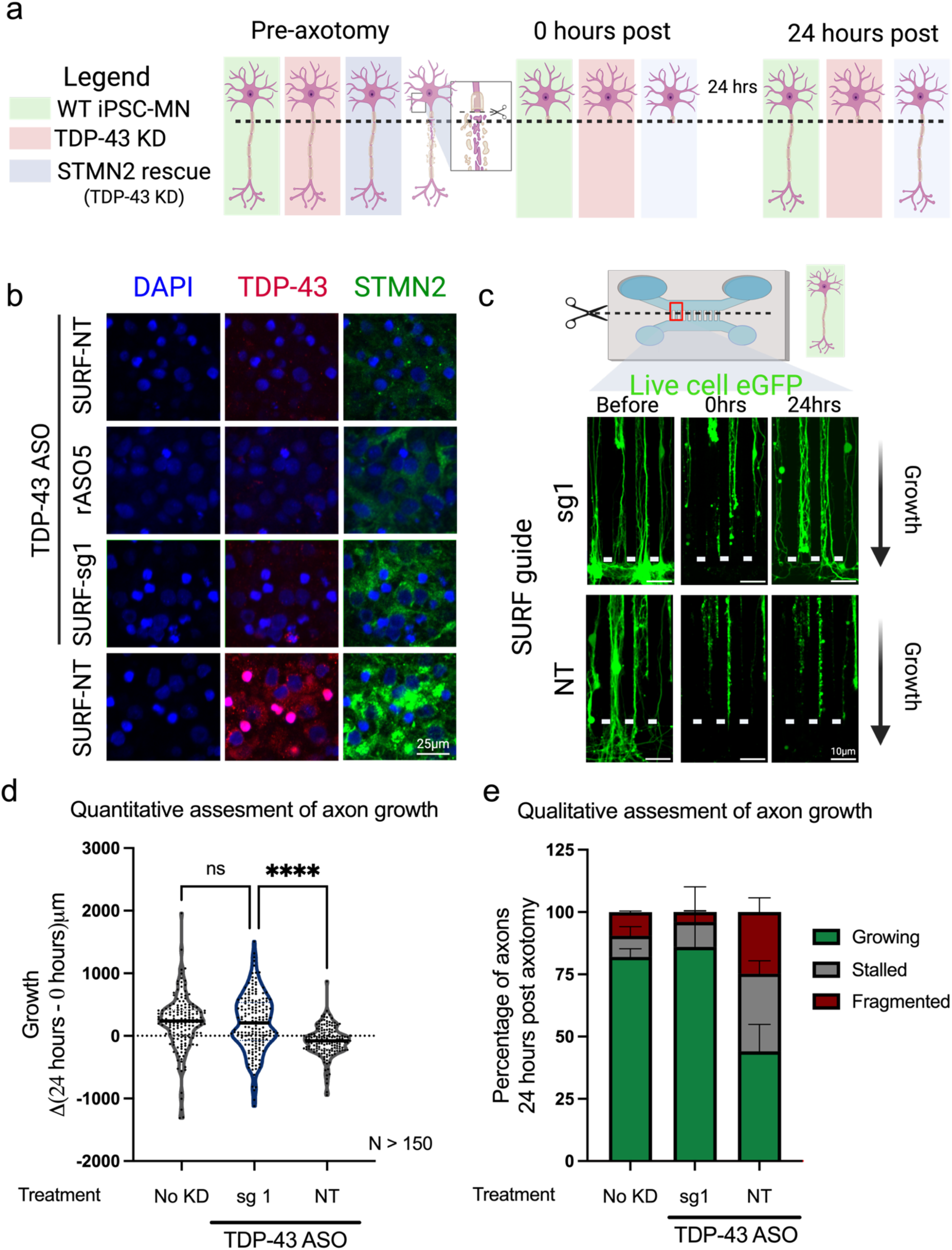
SURF-sg1 restores axon regeneration in iPSC derived motor neurons. a) Schematic representation of the axotomy study. b) Representative IF images of iPSC-MN in the somatic compartment of axotomy chambers 14 days post-axotomy. Cells were pretreated with TDP-43 ASO or NT ASO and transduced with lentiviral vectors expressing SURF-NT or SURF-sg1, as indicated. TDP-43 in red, STMN2 in green. Scale bar = 25 µm. c) Representative live-cell images of eGFP in iPSC-MN pretreated with TDP-43 ASO and transduced with lentiviral vectors expressing SURF-NT or SURF-sg1, as indicated, in axotomy chambers at the indicated timepoints. White line represents the microchannel separating the somatic (bottom) from the axonal (top) compartments. Scale bar = 10 µm. d) Truncated violin plot with smoothing quantifying the change in axon length following axotomy. Each data point represents an individual channel/microgroove. The median is shown by the black bar. N>150 channels across two independent differentiations. One-way ANOVA with Dunnett’s correction. ****P < 0.0001; ns, not significant. e) Stacked bar plot of the percentage of channels with axons growing, stalled (not growing), or fragmented.

Strikingly, axonal regeneration was completely restored in cells treated with SURF-sg1 (median growth in um: No KD – 238um, sg1 – 209um, NT – -80um; Fig. 3c,d). Not only that, fragmentation and blebbing of motor axons was observed in 25% of *TARDBP* knock-down conditions treated with control SURF-NT, but < 5% when treated with SURF-sg1 (Fig. 3c, e). We conclude based on these findings, that SURF-sg1 restores functional stathmin-2 protein and increases stathmin-2 protein levels for at least 24 days after lentiviral delivery.

### In-vitro therapeutic specificity of STMN2 targeting snRNA

After confirming the *in vitro* efficacy of SURF-sg1 at the RNA, protein, and functional level, we moved onto determine if there were any major direct or indirect effects of *STMN2* targeting SURF on iPSC-MN. We performed RNA sequencing (RNA-seq) on polyA selected RNA from cells treated with targeting SURF (sg1 or sg2) with cells treated with non-targeting SURF-NT. Differential expression was assessed to identify any genes with significant changes in expression (DESeq2) (P-value < 0.05; |LFC| > 1; Fig. 4a,b). Only 25 and 9 differentially expressed genes were detected in sg1 and sg2 respectively and not enriched in any gene ontologies relevant to neuronal health or RNA processing (sg1: 7-up, 18-down; sg2: 5-up, 4-down; Fig. 4c). Additionally, there was little change in genes related to axonal health, synaptic signaling, stress markers, apoptosis, or neuronal markers between any snRNA condition and the no snRNA lentivirus (Fig. 4d,e; S7b,c). With no evidence of impacted cell health or neuronal function, we moved on to identifying splicing off-targets.

**Figure 4:**
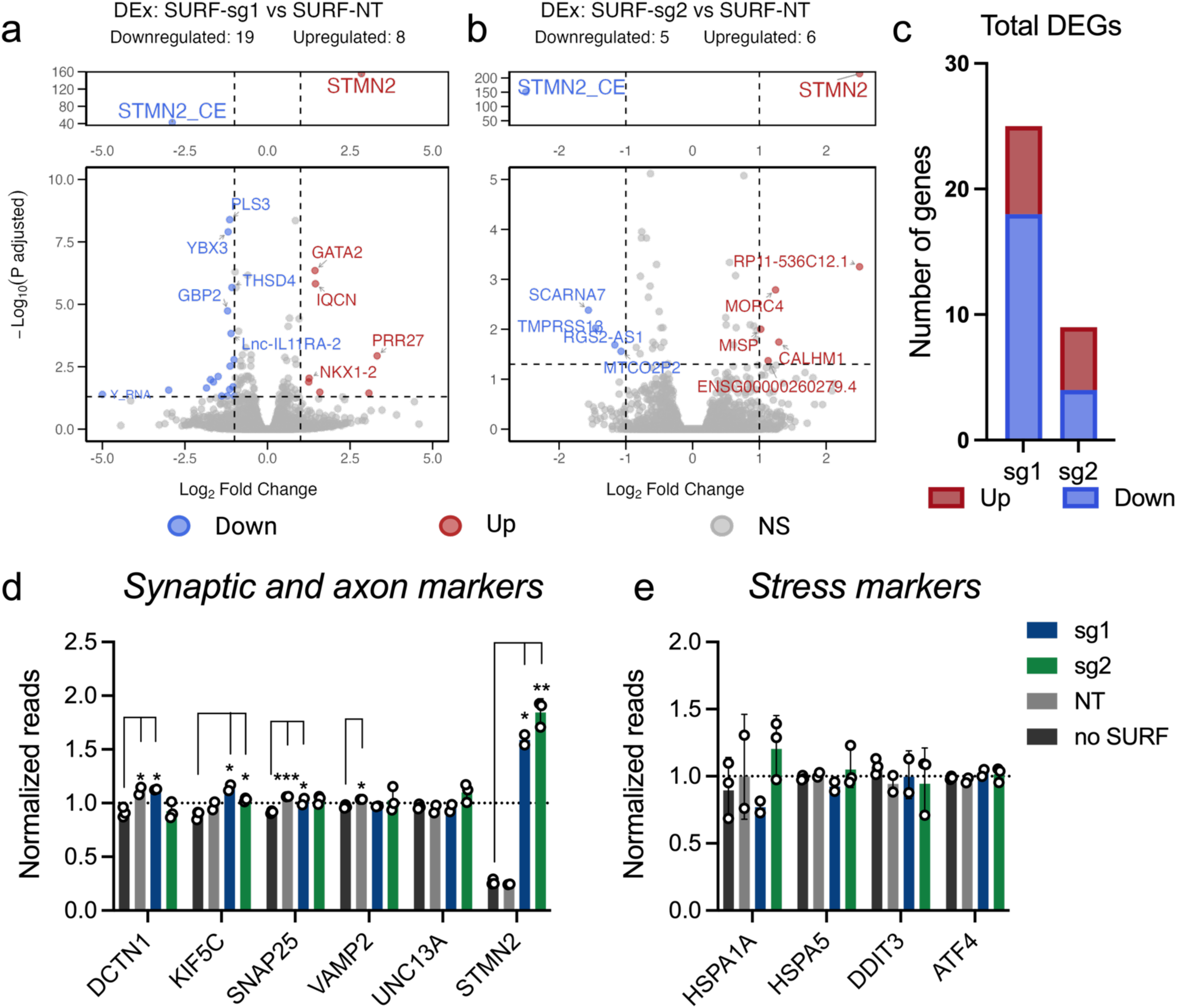
Off-target profile of SURF-sg1 and SURF-sg2 in iPSC derived motor neurons. a,b) Volcano plot of differential gene expression (DESeq2) comparing non-targeting (NT) to (a) SURF-sg1 or (b) SURF-sg2 snRNAs. Significantly differentially expressed genes (at threshold p<0.05 and >2-fold change) are indicated in blue (downregulated) and red (upregulated). NS, not significant (gray). c) Bar plot summarizing the number of differentially expressed genes (DEG) from (b) and (c). d,e) Bar plots showing normalized median of ratios RNA-seq read counts of canonical (d) synaptic and axon health, and (e) stress marker genes in iPSC-MN treated with TDP-43 targeting ASO and transduced with lentivirus expressing a SURF cassette (NT, sg1, sg2) or no SURF cassette (no SURF). N=3 biological replicates; sg2 and None; N=2 biological replicates; sg1 and NT. One-way ANOVA with Dunnett’s correction. ***P < 0.001; **P < 0.01; all others are not significant.

U7smOPT could potentially bind to non-target pre-mRNA and alter splicing in those genes. To identify any such events, we analyzed RNAseq data using the rMATS and MAJIQ packages to identify differentially spliced exons (P-value < 0.05; dPSI > 0.1) [57, 58]. Additionally, we used BLASTn to identify *in silico* any pre-mRNAs that SURF-sg1 or SURF-sg2 would have affinity for based on sequence homology and used the genes containing the top 500 hits by e-value in further analysis [59]. Only 2 splicing events overlapped between rMATS and MAJIQ results in the sg1 analysis, and neither was identified using BLASTn (Fig. S6a-c). These two events based on the RNA-seq analysis above did not have a significant impact on neuron health. Additionally, it is not possible to attribute them to direct snRNA binding due to the BLASTn results and thus could be due to experimental variation. Other events identified from two of the three methods were identified. These results demonstrate that SURF-sg1, like other U7smOPT snRNA vectors, is not only effective but has few off-targets of statistical significance in a disease relevant context, (2 alternative exons overlapping rMATs and MAJIQ out of 189 and 23 respectively)[40, 45, 60].

### In vivo studies

After demonstrating *in vitro* efficacy and safety, we tested the efficacy of SURF-sg1 *in vivo*. Recognizing that mouse *Stmn2* does not undergo cryptic splicing, we used a transgenic humanized mouse model where a 3.2 kb stretch of the human *STMN2* intron 1 containing the cryptic exon was introduced into the mouse *Stmn2* intron 1 locus. TDP-43 binding sites in the cryptic exon of this region were replaced with an MS2 hairpin, *Stmn2^HumΔGU^*, preventing the binding of mouse TDP-43 to the cryptic exon (Fig. 5a) [33]. This allele produces only truncated *Stmn2* mRNA, and virtually no full-length *Stmn2*. This is preferable over a TDP-43 sensitive model due to the inherent challenges of *in vivo* TDP-43 models [61].

**Figure 5:**
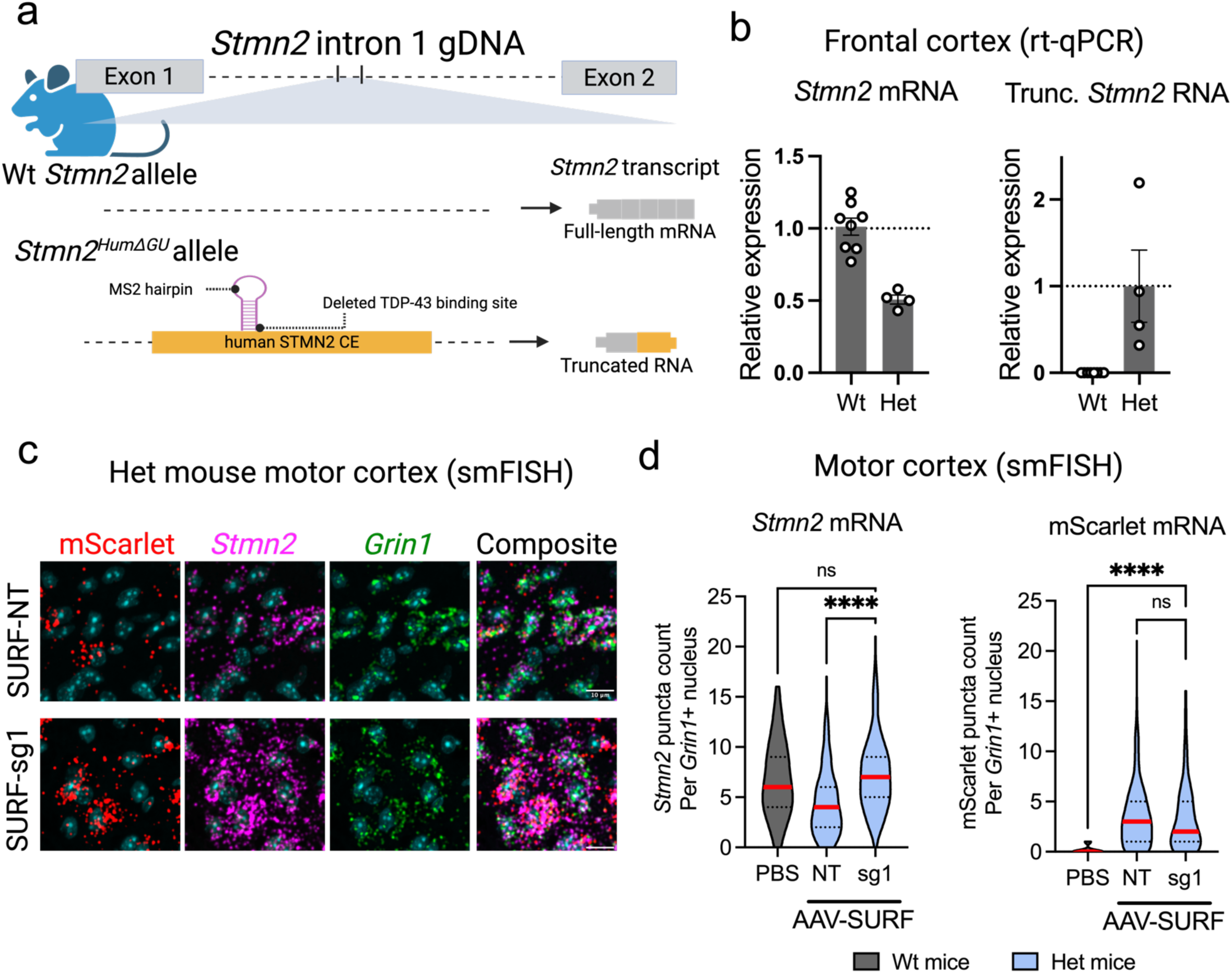
SURF-sg1 rescues *Stmn2* splicing in motor cortex of a humanized mouse model. a) Schematic depicting *Stmn2* intron 2 in the *Stmn2*^HumΔGU^ transgenic allele with the wild-type allele for reference. The transgenic allele produces only the cryptic exon. b) Bar plots quantifying *Stmn2* mRNA isoforms in cerebral cortex of *Stmn2*^HumΔGU/WT^ (Het) or *Stmn2*^WT/WT^ (Wt) mice treated with AAV9 delivering SURF-sg1. Data are rt-qPCR normalized to *Rps9* mRNA (ΔC_q_), relative to normalized levels in Het mice (trunc. *Stmn2* RNA) mice or Wt mice (full-length *Stmn2* mRNA) treated with non-targeting snRNA (ΔΔC_q_). RNA was extracted in bulk from the cortex of individual mice, with each datapoint representing one mouse. N=8; Wt mice; N=4; Het mice. c) Representative images of single-molecule FISH for mScarlet (red), *Stmn2* (pink), and *Grin1* (green) mRNAs in motor cortex of *Stmn2*^HumΔGU/WT^ mice. DAPI is in cyan. Scale bar = 10 µm. d) Violin plots quantifying the number of *Stmn2* mRNA puncta per *Grin1* mRNA-positive nucleus in mouse motor cortex. Each data point represents an individual nucleus. N=2 mice; NT and sg1 and > 250 nuclei; N=1 mouse for Wt-PBS; N > 60 nuclei. One-way ANOVA with Dunnett’s correction. ****P < 0.0001; ns, not significant.

For these studies, we focused on two mouse genotypes, *Stmn2^Wt/Wt^*(Wt) and the *Stmn2^HumΔGU/WT^* heterozygote (Het), which expressed 50% of the *Stmn2* mRNA as the Wt mouse (Fig. 5b). Mice were injected with self-complementary AAV9 (scAAV9) encoding SURF-sg1/NT and mScarlet (AAV-SURF-sg1/NT; 1e11 vg/mouse) at P0-P1 via intracerebroventricular injection (ICV). We selected the AAV9 capsid for its neurotropism and current use in the clinic [62]. Mouse brain tissue was harvested 30 days after injection.

We imaged individual mRNA molecules in the mouse motor cortex using single molecule RNA fluorescent *in-situ* hybridization (smFISH) to quantify three RNA species: *Stmn2* mRNA, mScarlet mRNA (AAV dissemination), and *Grin1* (ctrl) mRNA (Fig. 5c). After counting the puncta of each mRNA species for each nucleus across conditions (see methods), we filtered out nuclei where our positive control, *Grin1*, was not detected. From here, we quantified 60 nuclei for the wild-type mouse control and over 250 nuclei per condition for treated mice. We found robust rescue of *Stmn2* mRNA levels in the motor cortex of mice treated with AAV-SURF-sg1. Het mice motor cortices (Het MC) treated with AAV-SURF-NT had a mean of 4.42 *Stmn2* mRNA puncta per *Grin1*+ nuclei whereas Het mice treated with AAV-SURF-sg1 7.27 *Stmn2* puncta per nucleus, a 64% increase (p < 0.0001; N=2 mice; 175 nuclei per mouse; Fig. 5d left). Importantly, AAV-SURF-sg1 mice did not have statistically significant differences in *Stmn2* puncta compared to a wild-type mouse treated with PBS, suggesting we completely blocked cryptic splicing in the Het mice (p = 0.48; Wt-PBS – 6.75 *Stmn2* puncta/nucleus; Fig. 5d left). Additionally, AAV transduction (mScarlet) was consistent across AAV-SURF-sg1 and AAV-SURF-NT (Fig. 5d right)

### Suppression of *UNC13A* and *STMN2* cryptic splicing with a single vector

We took a similar approach to designing snRNAs to suppress cryptic splicing in *UNC13A* as for the *STMN2* cryptic exon, designing five guides to target both 3’ SS of the cryptic exon, and TDP-43 binding sites within the exon and intron (Fig. 6a). We tested these guides in an inducible *TARDBP* shRNA model in SH-SY5Y cells as the SH-SY5Y TDP-43^N352S/N352S^ cells we previously used (Fig. 1) and that do not express appreciable levels of the *UNC13A* cryptic isoform (data not shown).

**Figure 6:**
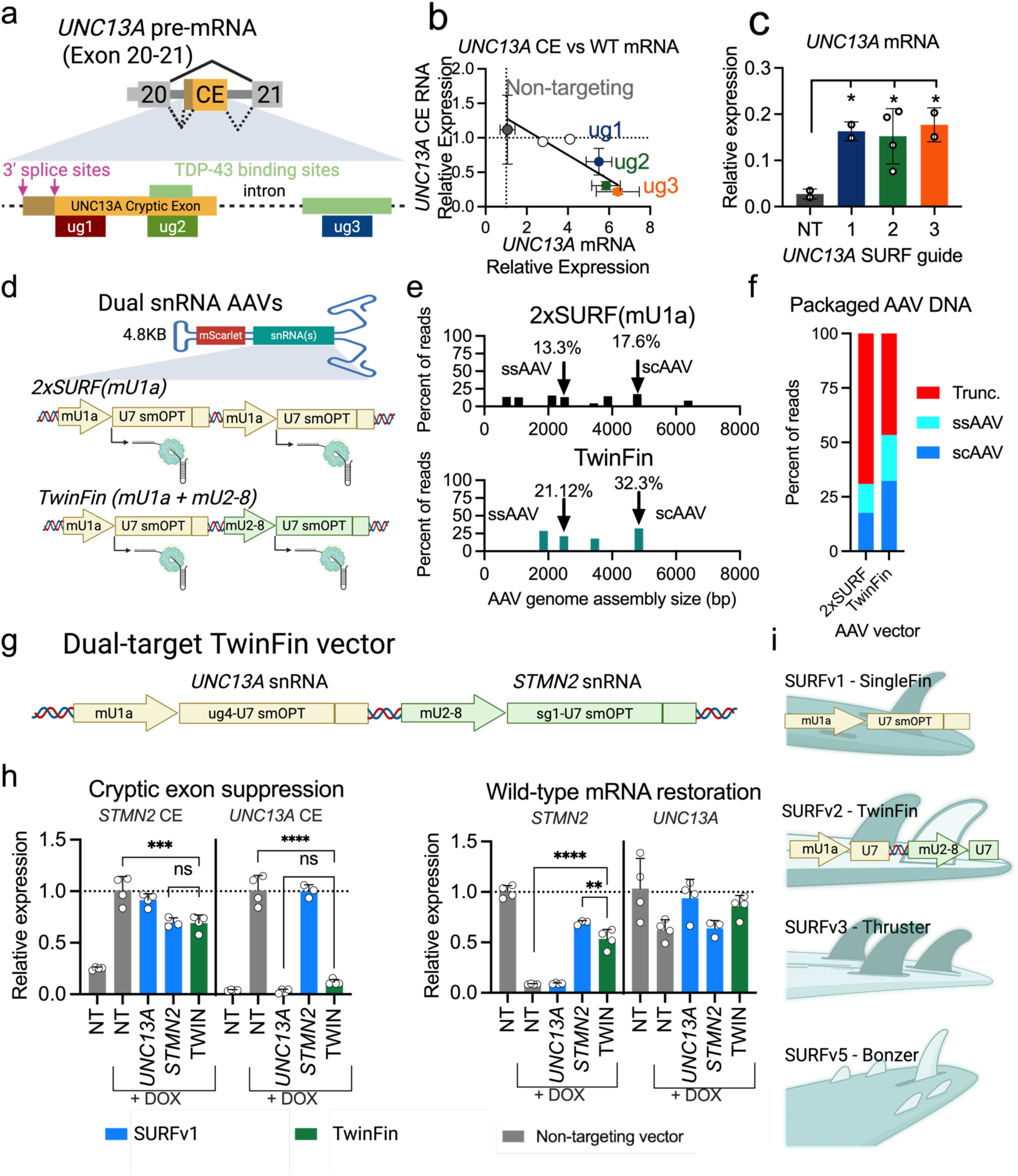
Optimized vector (*TwinFin*) simultaneously corrects cryptic *STMN2* and *UNC13A* splicing. a) Schematic illustrating the targeting strategy employed to block inclusion of the *UNC13A* cryptic exon. The 3’ splice sites and the first, TDP-43 binding sites from Brown et al 2022, and three 34 basepair *UNC13A* guide sites. b) Scatter plot showing levels of full-length and CE-containing *UNC13A* transcript isoforms in an induced stable SH-SY5Y cell line expressing a shRNA targeting *TARDBP* from a doxycycline-inducible promoter and transduced with lentiviral SURF vectors targeting the UG-rich regions. Data are rt-qPCR of the target RNA normalized to GAPDH (ΔC_q_), relative to normalized levels in cells transduced with non-targeting SURF vector. Error bars are only included for NT and ug1-3. N=2 biological replicates. c) Bar plot showing levels of the full-length *UNC13A* transcript normalized to GAPDH (ΔC_q_), relative to normalized levels in uninduced cells transduced with a non-targeting snRNA (ΔΔC_q_). N>=2biological replicates. One-way ANOVA with Dunnett’s correction. *P < 0.05. d) Schematic representation of two dual snRNA AAV vectors of ∼4.8kb containing either two mU1a cassettes (2xSURF) or an mU1a and an mU2-8 cassette (TwinFin). e,f) Distribution plots (e) and bar plot (f) showing the lengths of assembled genomes packaged into AAV9 for the two vectors in (d). Full-length single-stranded (ssAAV) and self-complementary (scAAV) genomes are indicated. g) Schematic of the dual-target TwinFin design. h) Bar plots of the levels of CE-containing (left) and full-length (right) *STMN2* and *UNC13A* transcript isoforms in SH-SY5Y transduced with an inducible lentiviral *TARDBP* shRNA induced with doxycycline and transduced with the lentiviral dual-target TwinFin vector (TWIN), the corresponding individual single-snRNA vectors, or a non-targeting (NT) snRNA. Data are rt-qPCR of target RNA normalized to GAPDH (ΔC_q_), relative to normalized levels in induced (left) or uninduced (right) cells transduced with the NT vector (ΔΔC_q_). N=3 biological replicates; STMN2 single vector; N=4 biological replicates; all other samples. One-way ANOVA with Dunnett’s correction. ****P < 0.0001; ***P < 0.001; **P < 0.01; not significant if not otherwise indicated. i) Schematic of the SURF and TwinFin vectors, with artistic representations of future vectors.

Interestingly, guides targeting TDP-43 binding sites suppressed the cryptic isoform more than the splice site targeting guides (∼50% reduction vs ∼75% reduction; ns; Fig. 6b). However, the wild-type *UNC13A* mRNA was restored to identical levels across splice site and TDP-43 binding site targeting guides (∼15% of full-length *STMN2*; p < 0.05 when compared to non-targeting; Fig. 6c). This is the same trend observed in targeting *STMN2* with sg1 and sg2(Fig. 1c,d; Fig s5b,c).

We next developed a dual-targeting vector. As we previously pointed out, regions of homology within AAV transgenes are reported to cause recombination, leading to AAV genome truncation. To test this, we produced scAAV9 with either 2 SURF cassettes (2xmU1a) or one mU1a cassette and one mU2-8 cassette (Fig. 6d). Both cassettes expressed sg1. We sequenced the packaged AAV genomes using long read sequencing, a quality control method in gene therapy to assess what percentage of AAV genomes are correctly packaged [63]. As expected, having one SURF and one mU2-8 cassette led to almost twice as much full-length (single-stranded and self-complementary) genome as with two tandem SURF cassettes (Fig 6e,f). Additionally, the tandem SURF cassettes led to eight separate assemblies, whereas the mixed cassettes led to only four (Table S1; Fig. 6e). We termed this new multi-valent snRNA vector *TwinFin*.

To validate *TwinFin* and ensure that these two cassettes are as efficient as either alone, we combined an *UNC13A* targeting guide (SURF-ug4) selected after preliminary dual-guide screening with sg1 expressed under the mU2-8 promoter (Fig. 6d; Fig. S7). We tested this vector in SH-SY5Y cells with inducible TDP-43 knockdown with snRNAs delivered by lentivirus. We found significant reduction of both the *UNC13A* cryptic exon and the *STMN2* cryptic exon with no significant difference between the single vector and dual vector (*UNC13A* CE: dual guide v NT control: 11.6% relative to NT control p < 0.0001; *UNC13A* CE; *STMN2* CE: dual guide v NT control: 69% relative to NT control p < 0.001; Fig. 6g).

## Discussion

In this work, we have identified U7smOPT as a viable vector for suppressing TDP-43 associated cryptic splicing. Further, we corrected *Stmn2* mis-splicing *in vivo* and optimized and developed a single viral vector for correcting mis-splicing in two pathogenic cryptic exons.

This work represents several novel strategies in the snRNA and gene therapy fields. Highlighting the wide recognition of *STMN2* and *UNC13A* as key ALS therapeutic targets, several biotechnology companies have entered pre-clinical development on one of these two mis-splicing events. While each of these studies is targeting one cryptic splicing event, our study enables the targeting of both events simultaneously with a single intervention. In the snRNA field, past studies have delivered four identical copies of a snRNA in a single AAV vector [45]. This strategy is risky due to the possibility of recombination due to homology between the snRNA cassettes. Our introduction of several promoters is particularly beneficial to the U7smOPT field as it increased potency enough to reduce the need for multiple copies in a single vector and simultaneously allows for targeting up to four targets without the risk of homology mediated recombination [64] [65]. Similarly, past efforts to use U7smOPT to suppress cryptic splicing have focused on events that result from DNA mutations. Our approach represents a departure from that, where we take an RBP centric approach and develop snRNAs that can recapitulate a specific RBP’s function [66]. Our *TwinFin* vector approach is novel in three ways: (1) it can target multiple mis-splicing events simultaneously, (2) it employs previously unused promoters to enhance potency, and (3) it corrects cryptic splicing events caused by splicing factor dysfunction, not DNA mutations.

There is a pressing need for disease modifying therapeutics for those with neurodegenerative disorders. Looking forward, iterations of this vector could be used in the clinic to treat additional TDP-43 proteinopathies in addition to ALS. Although causal relationships have yet to be established in humans, truncated *STMN2* containing the cryptic exon is detected in 82% of frontal cortex specimens from TDP-43 related frontotemporal dementia (FTD-TDP) patients, ∼80% of Alzheimer disease (AD) with TDP-43 inclusions (approximately 40% of AD cases), and is a hallmark of Limbic-predominant age-related TDP-43 encephalopathy (LATE-NC). A single dose of AAV could improve prognoses for each of these disorders.

This therapeutic is especially promising with the recent advances in AAV therapies and new platform based regulatory approaches taken by the FDA. As of 2024, 238 AAV-based therapeutics have entered clinical trials and seven AAV therapies have been approved, including two for CNS disorders [67]. Our snRNA therapeutic has strong potential to provide therapeutic benefit with a single-dose injection such as IV, IT, ICM, or subpial using AAV9.

## Methods

### RNA Extraction, Reverse Transcription, and Quantitative PCR

SH-SY5Y, HEK293T, and iPSC-derived motor neuron cultures were washed with PBS and lysed directly in RLT buffer (Qiagen RNeasy kit). Snap-frozen tissue samples were mechanically homogenized in 1 mL of RLT buffer using a glass douncer, and RNA was purified according to the manufacturer’s instructions. RNA concentrations were determined using a NanoDrop spectrophotometer (Thermo Scientific). Between 100–500 ng of total RNA was reverse-transcribed using the ProtoScript II First Strand cDNA Synthesis Kit (New England Biolabs) with oligo-dT primers. For rt-qPCR, resulting cDNA was diluted to 1–2 ng/μL, and 2–10 ng of cDNA was used per 10 μL reaction (Bio-Rad). qPCR was performed on a Bio-Rad C1000 thermal cycler with a 384-well format, and each condition was run in technical triplicate. Expression was normalized to one housekeeping gene (mouse *Rps9* or human *GAPDH*), and relative levels were calculated using the ΔΔCq method. Data visualization and statistical analysis were performed in GraphPad Prism 8.

For quantitative rt-PCR on snRNA targets, a protocol was adapted from Nia et al 2015 (Niu et al. 2015). RNA was first poly-adenylated using E. coli poly(A) polymerase (NEB), then reverse transcribed with an oligo-dT primer with a universal anchor. This cDNA was then used for qPCR with a universal reverse primer antisense to the anchor and targeted forward primers.

### Immunoblotting

Protein lysates were prepared from cultured cells and tissue samples using RIPA buffer, and concentrations were equalized via Bradford assay (Bio-Rad). Samples were denatured in SDS loading buffer and run on polyacrylamide gels before transfer to 0.2 μm nitrocellulose membranes. Membranes were blocked in 5% milk/TBST for 1 hour, then incubated overnight with primary antibodies including anti-Stathmin-2 (Novus, NBP1-49461). Following washes in TBST, membranes were probed with HRP-conjugated secondary antibodies (1:5,000–1:10,000) in 5% milk for 1 hour at room temperature. Detection was performed via chemiluminescence on film or a LICOR Fc imaging system.

### Non-iPSC cell Culture

HEK293T (LentiX Takara), SH-SY5Y (ATCC CRL-2266), and U2OS cells (ATCC HTB-96) were maintained in DMEM (Gibco) supplemented with 10% fetal bovine serum (Omega) at 37°C in a humidified 5% CO₂ incubator. TrypLE was used for dissociating cells and passaging, which was done every 3-5 days.

### Transient transfection

HEK293T (Takara) cells were seeded 24 hours prior to transfection in a 24-well plate. Cells were transfected using Jet Optimus transfection reagent according to manufacturer’s protocol. 48 hours post-transfection, cells were assessed for eGFP expression and harvested for RNA levels using identical RNA extraction protocols as above.

### Lentivirus production

Custom snRNA-expressing constructs were designed using Benchling and synthesized as gene fragments (Twist Bioscience) or oligonucleotides (IDT). These were assembled into lentiviral transfer vectors using Gibson assembly or site-directed mutagenesis. Lentiviral particles were produced via standard Gen2 three-plasmid transfection into LentiX packaging cells (Takara) using Trans-IT VirusGen reagent (Mirus). Virus was collected 72 hours post-transfection, filtered, concentrated using either sucrose ultracentrifugation or LentiX Concentrator (Takara), and stored at –80°C. For inducible shRNA experiments, doxycycline was added concurrently with viral transduction, and cells were collected six days later for downstream analysis. For experiments in iPSC-MN, virus was functionally titered on HEK293T cells and titers were adjusted before transduction.

### iPSC Motor Neuron Differentiation

A previously characterized iPSC line (CV-B, Coriell GM25430; ref: PMID 21368825, 24239350) was maintained on Matrigel in mTeSR-Plus medium. Neural induction was initiated at 70–90% confluency by switching to N2B27 medium (DMEM/F12 + GlutaMAX, N2 and B27 supplements, ascorbic acid, SB431542, dorsomorphin, CHIR99021). After 7 days, cells were passaged 1:6 and switched to N2B27 medium with SB431542, dorsomorphin, Smoothened agonist (SAG), and retinoic acid (RA). On day 18, cells were replated onto Matrigel-coated 10 cm dishes and cultured in SAG/RA-containing media. Final plating into microfluidic devices, 24 well, or 12 well plate occurred on day 25 using Accutase, with chambers pre-coated in poly-D-lysine, poly-L-ornithine, and laminin. From day 25 onward, cells were maintained in N2B27 with neurotrophic factors (BDNF, CTNF, GDNF) and DAPT, with media refreshed every 2–3 days. For experiments using an ASO for TDP-43 knock-down, the ASO was delivered with a full media change at day 30 and maintained during half media changes every three days. Any lentivirus treatments were performed at day 32.

### Axotomy and Microfluidic Culture

Custom microfluidic chambers were seeded with day-25 iPSC-MN precursors and matured for five days before treatment, as stated above. At day 42, axons in the distal compartment were removed via aspiration-induced axotomy. Axonal regrowth was assessed at before, 0, 24, 48, and 72 hours after axotomy. Samples were fixed in 4% paraformaldehyde and processed for immunofluorescence with anti-Stathmin-2 (1:1000, Novus). Alexa Fluor-conjugated secondary antibodies were used (Jackson ImmunoResearch), and immunofluorescent confocal imaging was performed on an Andor Dragonfly microscope.

iPSC-MN cells were treated with either TDP-43 ASO or non-targeting ASO maintained at 10 µM from day 30 to 42, with half-media changes performed every 3 days. Lentivirus was delivered at day 32 and contained an eGFP-PuroR expression cassette.

### Skeletonization and Quantitative Analysis of Axonal Regrowth

We developed an analysis pipeline to quantify axon growth in individual channels. This pipeline takes in eGFP images and user inputted coordinates of individual axon channels. Then, a binary mask is created for each microgroove channel. A skeleton image was generated from this binary mask using the skimage tool of the scikit-image library [68]. Skeletonization reduces neurites to centerlines while preserving topology, enabling quantification of total axonal length. The total skeleton length was quantified in individual microgrooves and the length of the axons at 0 hours was subtracted from the length at 24 hours to calculate the amount of growth 24 hours post axotomy (Fig. 4c). Axons were quantified across two chambers for each condition, and N > 150 channels total. Further qualitative analysis was performed to investigate axonal health after 24 hours using a standardized rubric (Fig. 4d).

### Statistical Analysis and Compliance

All statistical tests were performed using GraphPad Prism 8. Biological replicates represent independent differentiations, animal samples, or wells treated independently. Two-tailed Student’s t-tests were used for two-group comparisons; one-way ANOVA Dunnett’s correction were used for comparisons involving more groups unless otherwise stated. Significance thresholds were set as follows: ****P < 0.0001; ***P < 0.001; **P < 0.01; *P < 0.05; n.s., not significant. Unless otherwise stated, error bars represent standard error of the mean. All RNA-seq gene expression data was normalized using the median of ratios methods in the DESeq2 package. Replicates that failed due to technical errors (e.g., inconsistent signal unrelated to guide activity) were excluded from analysis. These exclusions were rare and did not affect overall conclusions. In guide screening in Figure 1, two guides were removed where the standard deviation of the cryptic exon expression > 0.3.

### RNA-seq and Computational Analysis

Total RNA was isolated as described above. For transcriptome profiling, poly(A)-selected RNA was used to prepare stranded mRNA libraries following the manufacturer’s protocol, then sequenced on an Illumina platform to obtain 100 base pair paired-end reads. Samples used in differential expression and splicing analysis were sequenced to a minimum depth of 8e7 paired end reads. Gene-level counts were generated against a current human reference annotation; downstream normalization used the DESeq2 median-of-ratios method.

We compared SURF-sg1 or SURF-sg2 to SURF-NT controls. DE analysis was performed in DESeq2 with default dispersion estimation and Wald tests. Genes were called differentially expressed at P < 0.05 and |log2 fold change| > 1, as reported in the main text/legends. Summary volcano plots and DEG counts correspond to Fig. 4a–c.

Marker panels for synaptic/axonal health, stress, apoptosis, and neuronal identity were summarized from normalized counts (Fig. 4d–e; Fig. S7). No gene-ontology categories relevant to neuronal health or RNA processing were enriched among DEGs under the thresholds above. To assess unintended splice changes, we analyzed the same RNA-seq datasets with rMATS and MAJIQ using default event models. Events were considered significant at P < 0.05 and ΔPSI (|dPSI|) > 0.1. Concordant events between tools were rare (2 overlaps for sg1) and did not map to high-homology targets (see below), consistent with minimal off-target splicing (Fig. S6).

To contextualize potential direct binding off-targets, guide sequences were searched by BLASTn against human transcript models, and the top 500 hits by E-value were carried forward as a candidate gene set for intersection with the splicing results.

### Mouse studies

Cohorts of progeny between a *Stmn2*^HumΔGU/WT^ and *Stmn2*^WT/WT^ mice were injected during the first 24 hours after birth with 1e11 scAAV9 expressing an mScarlet fluorescent reporter and either a targeting or non-targeting U7smOPT driven by an mU1a promoter. After 30 days, the mouse central nervous systems were harvested. Harvested mouse brains were divided in half, with the cortex of one half going towards rt-qPCR of full-length and truncated *Stmn2*. The second half was fixed in 4% PFA for 24 hours at 4C, then moved sucrose. The brain hemispheres were then frozen in OCT and stored at -80 C. Mouse hemispheres used for smFISH were sectioned using a cryostat into 25 µm sagittal slices. Slides were then stained for *Stmn2*, *Grin1* and mScarlet mRNA using ACD RNAScope probes.

### Nuclei Segmentation and Quantification of STMN2 and mScarlet Co-expression (editing)

Nuclei were segmented from DAPI-stained images using Cellpose with the pre-trained nuclei model and a diameter set manually to match the average nuclear size observed in the dataset. Images were processed in grayscale, and Cellpose-generated masks were post-processed to remove small artifacts using morphological opening and minimum size filtering (area > 28 µm²)[69].

To detect individual RNA puncta for *Stmn2*, mScarlet, and *Grin1*, a watershed-based segmentation strategy was applied independently to each gene channel. Puncta were detected by applying adaptive Otsu thresholding (scaled by a factor of 0.75 per gene). Distance transforms were calculated on the thresholded binary masks, and local maxima were identified using `peak_local_max` with a 3×3 footprint to generate watershed seeds. Watershed segmentation was then performed on the inverted distance map, and small segmented objects (<3 pixels) were removed to minimize false positives.

For each nucleus, the number of *Stmn2*, mScarlet, and *Grin1* puncta was quantified by overlaying the nucleus mask onto the respective puncta segmentation masks. Only nuclei that had more than one *Grin1* mRNA puncta (used as a quality filter) were included in subsequent analysis.

## Ethics approvals

All procedures involving animals and human stem cells were approved by UCSD IACUC protocol S0022, UCSD IRB protocol #161575ZX, and Jackson Laboratory AUS protocol 20029.

## Acknowledgements

We are grateful to Cody Fine and Mitra Banihassan (UCSD) for technical assistance with flow cytometry experiments. This work was supported by the Human Embryonic Stem Cell Core Facility and the Genomics Core Facility of the Sanford Consortium of Regenerative Medicine GT3 Core Facility of the Salk Institute with funding from NIH-NCI CCSG: P30 CA014195, by the UC San Diego Stem Cell Program, and a CIRM Major Facilities grant (FA1-00607) to the Sanford Consortium for Regenerative Medicine. This work was funded by a UCSD Gene Therapy Initiative Seed Grant and NIH grants R01 NS103172 and HG004659 to G.W.Y. T.A.G was supported in part by National Institute of General Medical Sciences Predoctoral Basic Biomedical Sciences Research Training Program T32 GM145427. A.A.S. was supported by a Biomedical Research Fellowship from the Hartwell Foundation.

## Competing interests

G.W.Y. is a cofounder, member of the Board of Directors, Scientific Advisory Board member, equity holder, and paid consultant for Eclipse BioInnovations. G.W.Y.’s interests have been reviewed and approved by the University of California San Diego, in accordance with its conflict-of-interest policies. All other authors declare no competing interests.

## Figures and Tables

**Figure S1:**
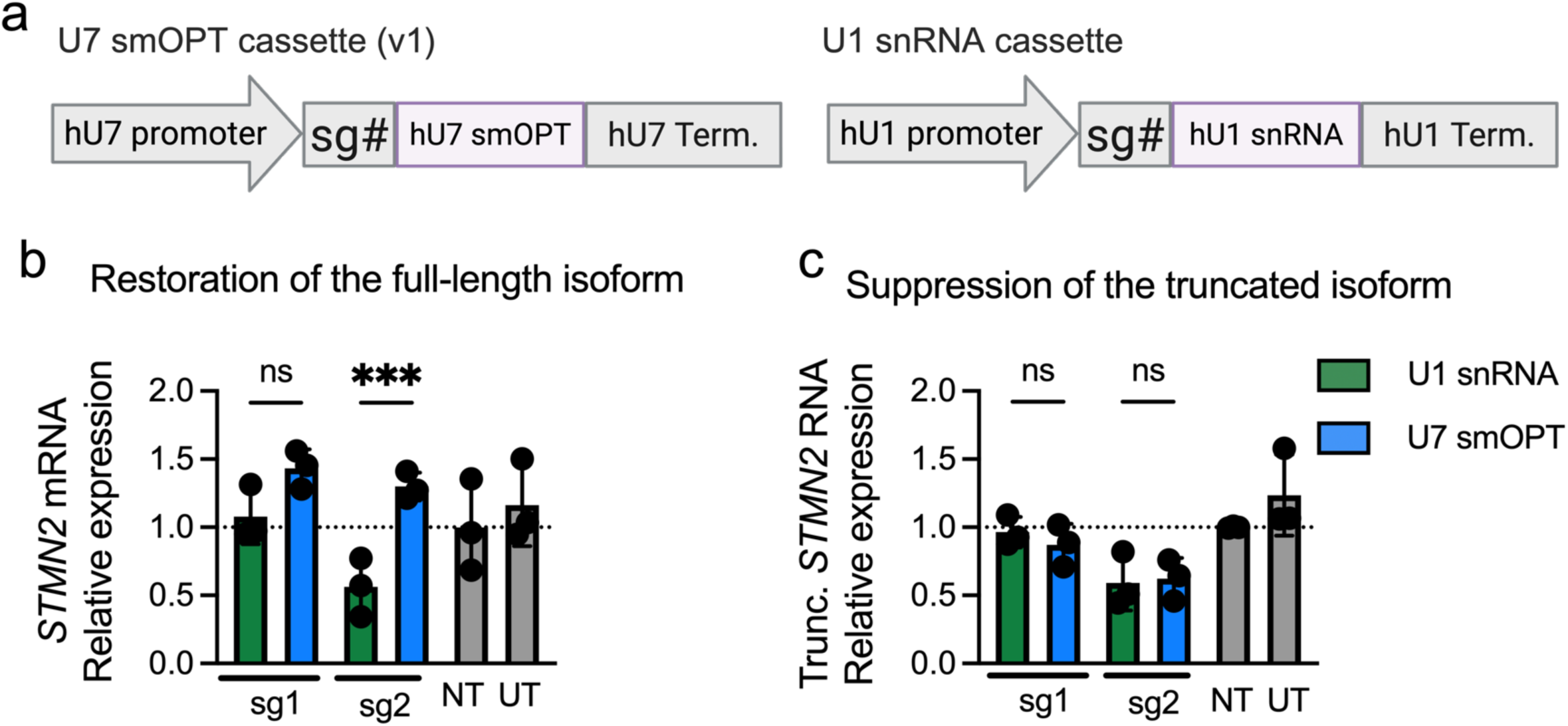
U1 snRNAs suppress cryptic exon without increasing full-length STMN2. a) Schematic of U7smOPT and U1 snRNA expression cassettes. b,c) Bar plots showing levels of (b) the full-length *STMN2* isoform or (c) the truncated *STMN2* isoform in SH-SY5Y^N352S/N352S^ cells after treatment with lentivirus expressing U7smOPT or U1 snRNAs targeting exon 2a. The NT snRNA is an U1 snRNA with a 15 base pair non-targeting sequence. N=3 biological replicates. One-way ANOVA with Sidak correction. ***P < 0.001; ns, not significant.

**Fig S2:**
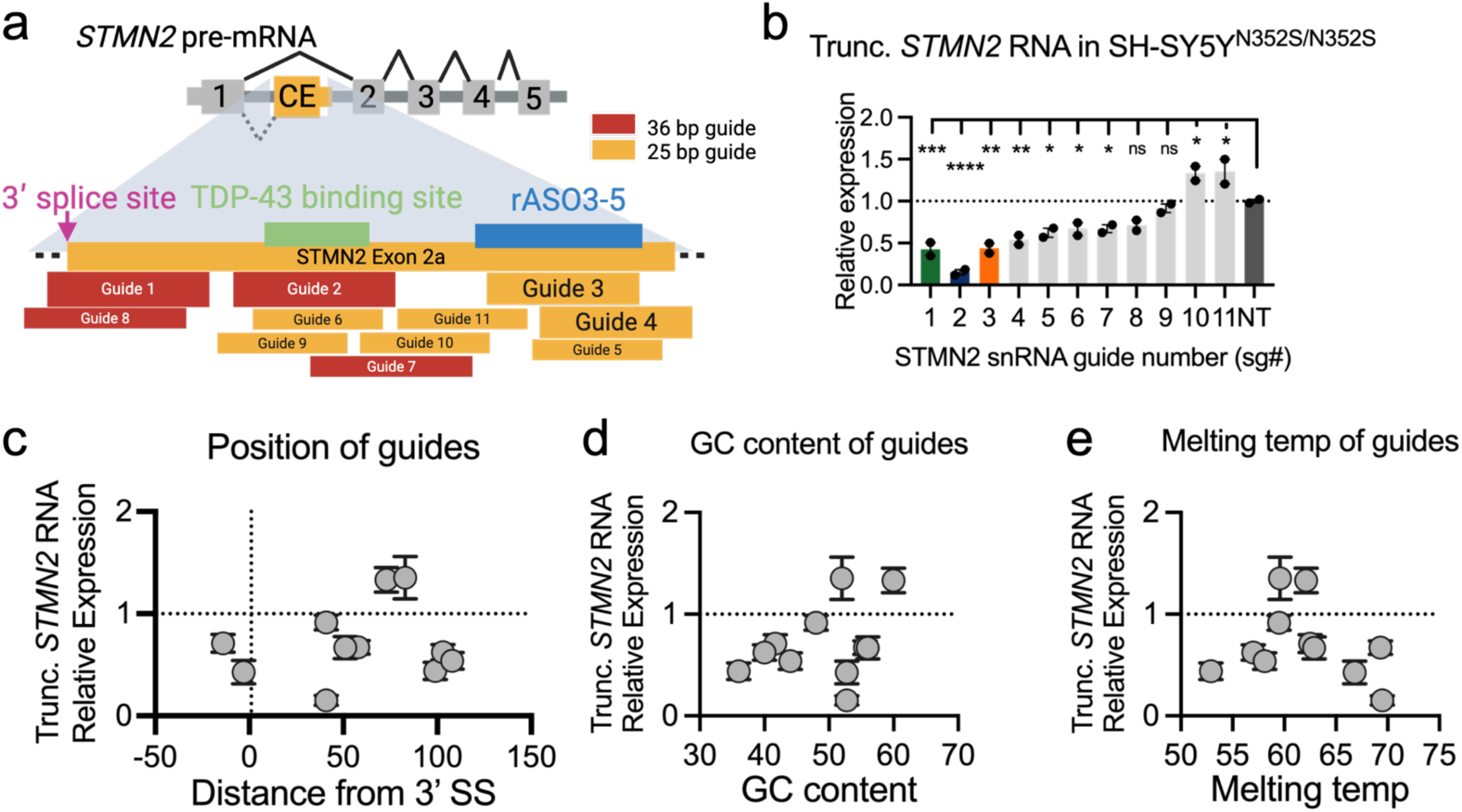
snRNAs of various qualities suppress STMN2 cryptic splicing. a) Schematic representation of the *STMN2* cryptic exon and the location of 10 snRNA targets. TDP-43 binding sites and the targeting site of rASO3-5 from Baughn et al., 2023 are indicated. Guides of 25 (orange) or 36 base pairs (red) were tiled along the CE region. b) Bar plot showing levels of the truncated *STMN2* isoform in SH-SY5Y^N352S/N352S^ cells transduced with U7smOPT expressing lentivirus targeting the *STMN2* cryptic exon region. Data are rt-qPCR of truncated *STMN2* mRNA normalized to GAPDH (ΔC_q_), relative to normalized levels in cells transduced with non-targeting (NT) smOPT (ΔΔC_q_). N=2 biological replicates. One-way ANOVA with Dunnett’s correction. ****P < 0.0001; ***P < 0.001; **P < 0.01; *P < 0.05; ns, not significant. c-e) Scatter plots of levels of the truncated *STMN2* isoform in SH-SY5Y^N352S/N352S^ cells following lentiviral delivery of the 10 snRNA guides, based on their (c) distance from the 3’ splice site, (d) GC content, and (e) predicted melting temperature. Each point represents N=2 biological replicates.

**Figure S3:**
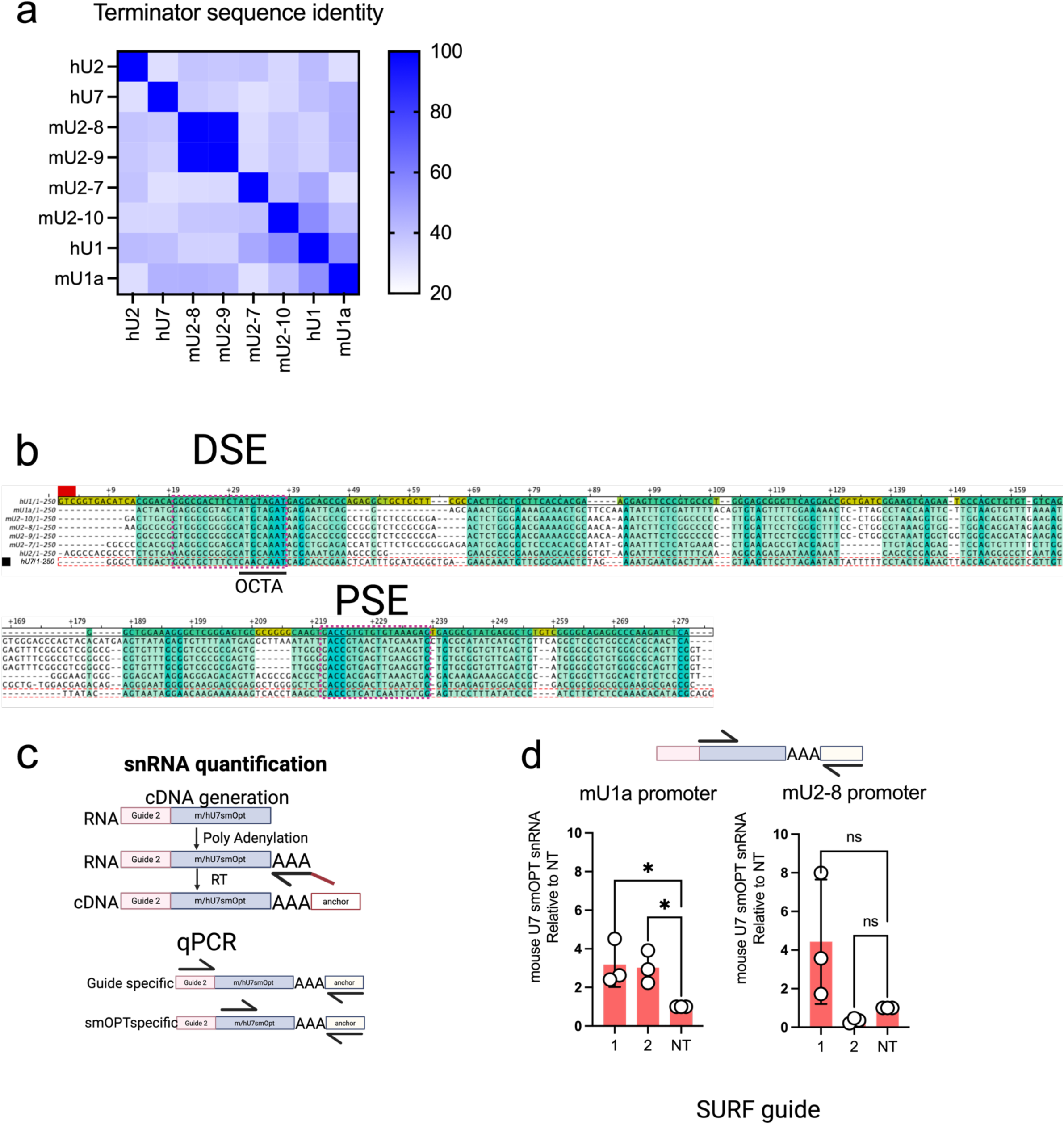
Extended data on novel promoters. a) Heatmap showing percent nucleotide identity of the terminators used in this study. b) Multiple sequence alignment of the promoters used in this study. Nucleotides are shaded by degree of conservation across sequences. The distal sequence element (DSE), proximal sequence element (PSE), and OCTA binding site are indicated. c) Schematic of polyadenylation and reverse transcription used to generate cDNA for snRNA quantification for rt-qPCR. d) Bar plots showing levels of snRNAs in iPSC-MN by rt-qPCR with forward primer against the snRNA backbone sequence, rather than the guide region used in Figure 2. Data are rt-qPCR of target RNA normalized to PuroR transcript levels (ΔCq), relative to normalized levels in cells transduced with non-targeting (NT) snRNA (ΔΔCq). One-way ANOVA with Dunnett’s correction. N=3 biological replicates. *P < 0.05; ns, not significant.

**Figure S4:**
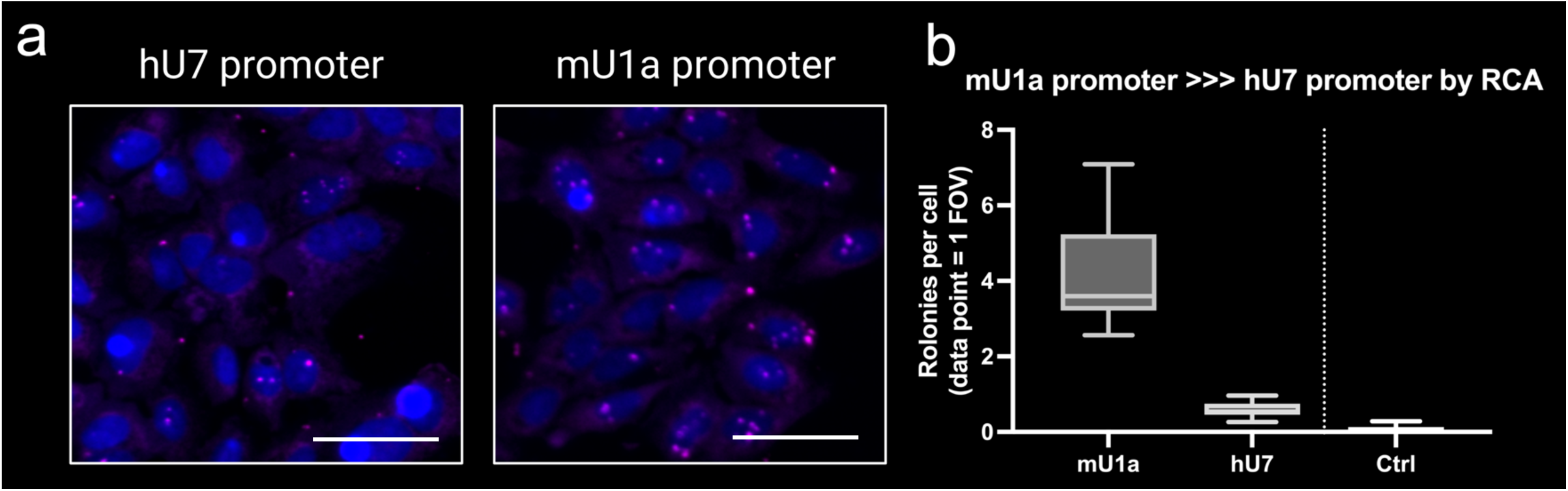
Rolling circle amplification detection of guide 2 sequence. a) U2OS cells were transduced at low MOI with lentivirus expressing human U7smOPT with sg1 from the hU7 promoter or mouse U7smOPT with sg1 from the mU1a promoter. Scale bar = 40 µm. b) Box plot showing the quantification of rolonies per field of view (FOV; N>7).

**Figure S5:**
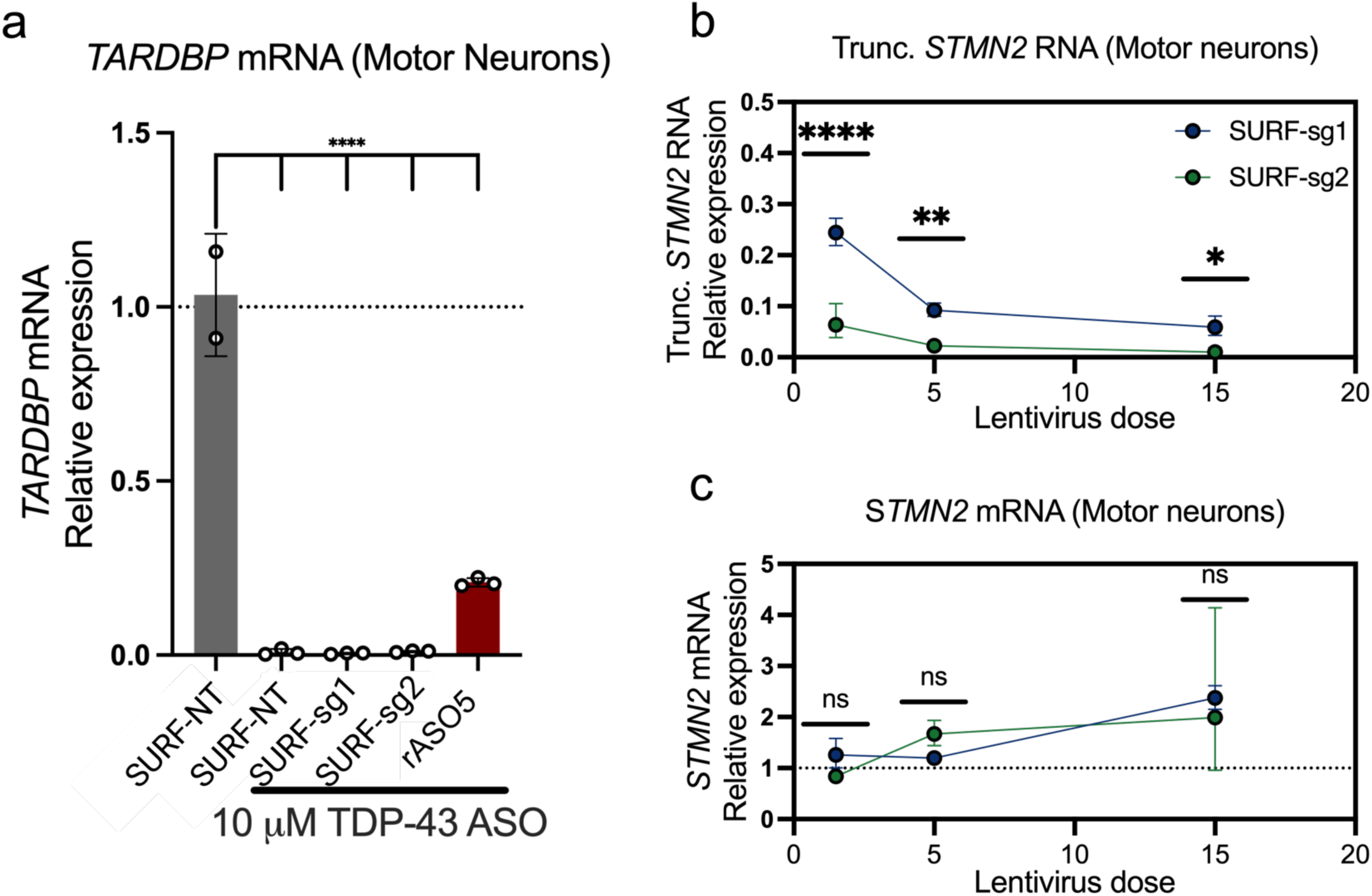
Extended data on suppression of cryptic splicing in iPSC derived motor neurons. a) Bar plot of *TARDBP* transcript levels in iPSC-MN transduced with lentiviral vectors expressing the indicated snRNAs from a SURF cassette or treated with rASO5, following treatment of cells with either a *TARDBP* mRNA-targeting ASO or a non-targeting (NT) control. Data are rt-qPCR of *TARDBP* mRNA normalized to *GAPDH* mRNA, relative to normalized *TARDBP* mRNA levels in cells treated with NT ASO and transduced with SURF-NT expressing lentivirus. Lentiviral vector doses are the mid-level doses shown in Fig. 2f,g. N=2 biological replicates; SURF-NT; N=3 biological replicates; all other samples. One-way ANOVA with Dunnett’s correction. ****P < 0.0001. b,c) Line plot showing levels of truncated (b) and full-length (c) *STMN2* mRNA in iPSC-MN transduced with increasing relative doses of snRNA expressing lentivirus following ASO-mediated TDP-43 depletion. Data are rt-qPCR of target RNA normalized to GAPDH mRNA, relative to normalized levels in cells treated with (b) TDP-43 ASO or (c) control ASO and transduced with lentivirus expressing SURF-NT. N=3 biological replicates. One-way ANOVA with Dunnett’s correction. ***P < 0.001; **P < 0.01; *P < 0.05; ns, not significant.

**Figure S6:**
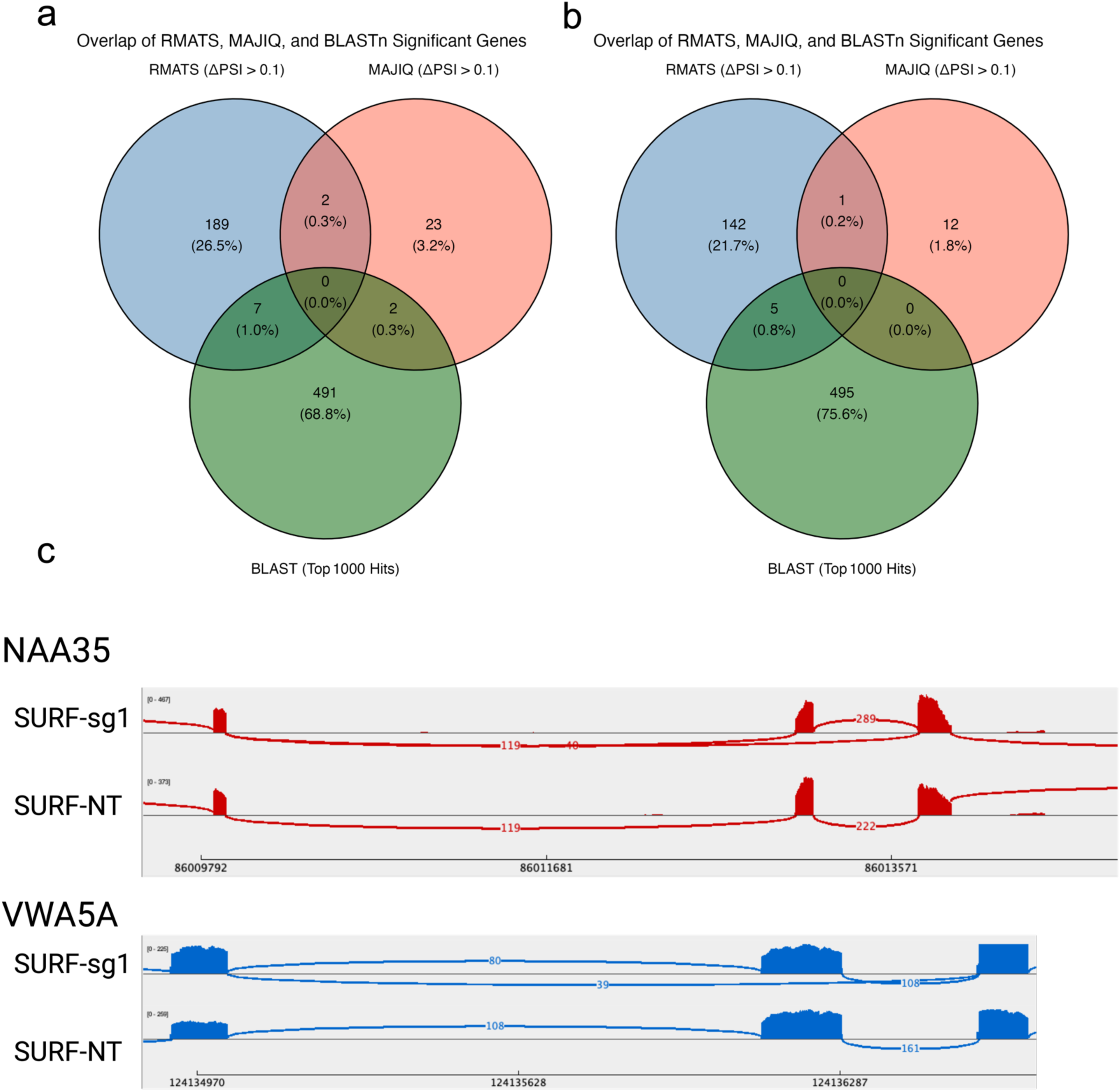
Characterization of off-target splicing events. a,b) Venn diagrams of exon skipping events detected by rMATS, MAJIQ, and genes with regions of homology to the *STMN2* cryptic-exon targeting guide for SURF-sg1(left) and SURF-sg2 (right) compared to SURF-NT. c) Genome browser views of skipped exons in *NAA35* (top) and *VWA5A* (bottom) identified for SURF-sg1 and SURF-NT.

**Figure S7:**
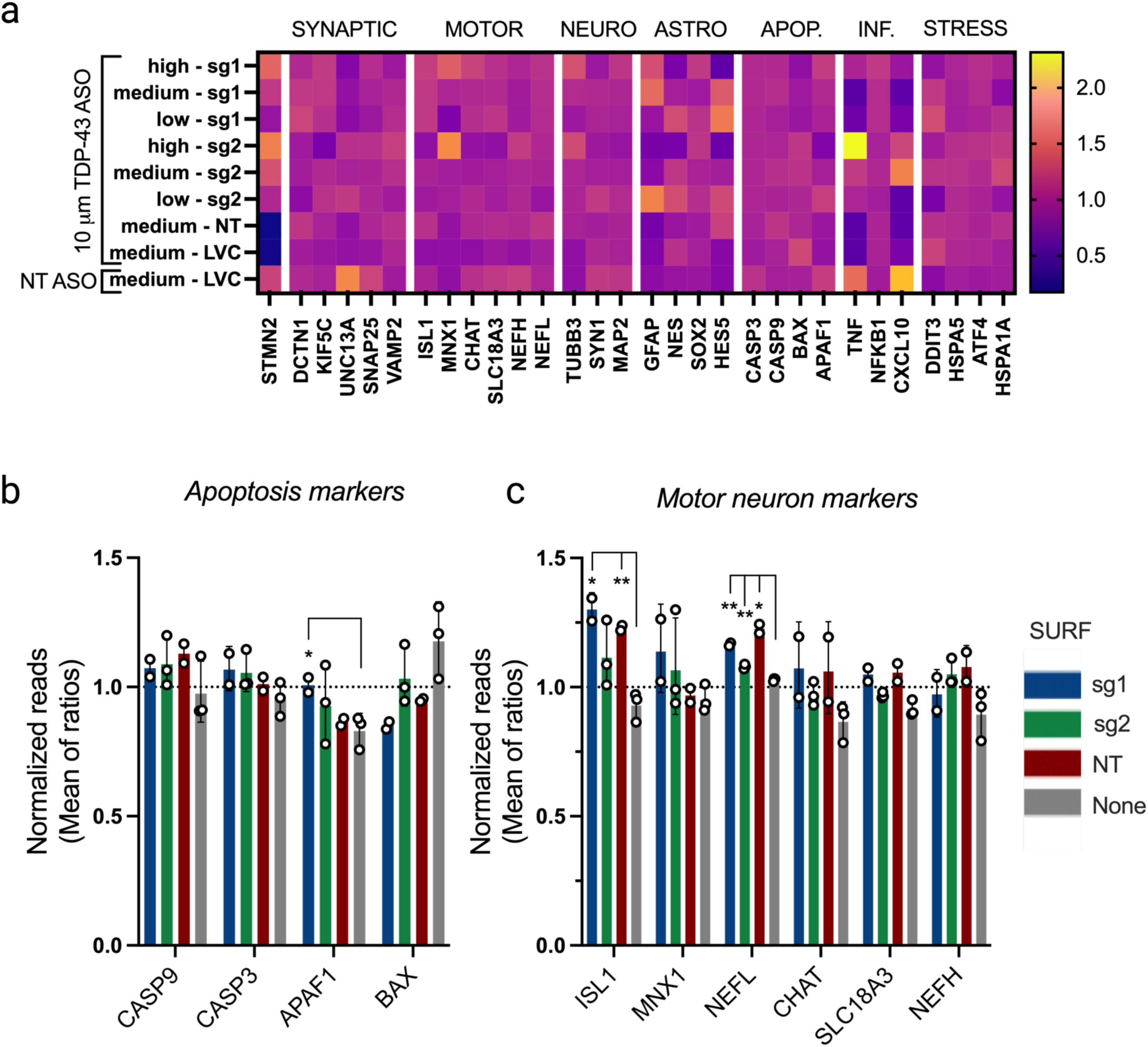
Expanded summary of RNA expression of genes relevant to neuronal health in iPSC-MN treated with SURF vectors. a) Heatmap showing marker gene transcript levels in iPSC-MN transduced with either a high, medium, or low (15, 5, or 1 relative transduction units) dose of lentivirus expressing SURF-sg1, SURF-sg2 or SURF-NT with TDP-43 ASO. LVC indicates lentivirus control. Data are RNA-seq median of ratios methods (DESeq2) normalized to the average mean of normalized reads across all samples. b,c) Levels of apoptosis marker (b) and motor neuron marker (c) transcripts in iPSC-MN transduced with sg1, sg2, or non-targeting (NT) sgRNAs, or in untransduced cells. Data are RNA-seq median of ratios methods (DESeq2) normalized to the average across all samples. N=3 biological replicates; sg2 and None; N=2 biological replicates; sg1 and NT. One-way ANOVA with Dunnett’s correction. **P < 0.01; *P < 0.05; not significant unless otherwise indicated.

**Figure S8:**
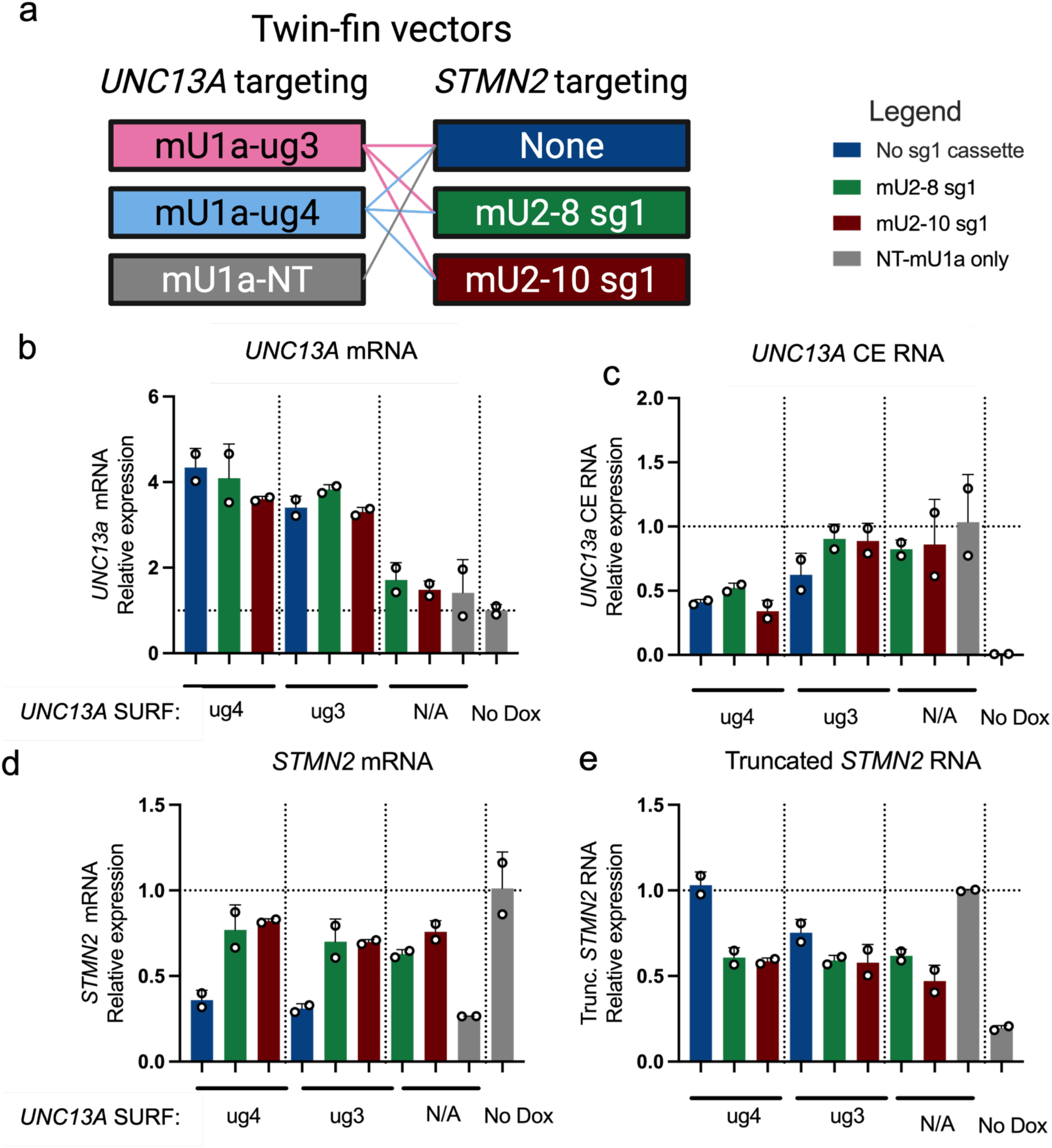
Results of screening ug3 and ug4 with sg1 in a single vector. a) Diagram depicting the combinations of snRNA cassettes screened here, including ug3- and ug4-targeting designs under the mU1a promoter tested with sg1 expressed under mU2-8 or mU2-10. b-e) Bar plots showing levels of (b) full length *UNC13A*, (c) CE-containing *UNC13A*, (d) full-length *STMN2*, and (e) truncated *STMN2* isoforms in a stable SH-SY5Y cell line expressing a shRNA targeting *TARDBP* from a doxycycline-inducible promoter and induced or undinduced cells (No Dox). Cells were treated with either a single snRNA vector or dual snRNA vector targeting one or both transcripts, or a non-targeting vector, as indicated. Data are rt-qPCR of target RNA normalized to GAPDH (ΔC_q_), relative to normalized levels in cells transduced with and non-targeting snRNA (ΔΔC_q_). For full-length transcripts, ΔΔC_q_ was calculated using the uninduced control. For mis-spliced transcripts, ΔΔC_q_ was calculated relative to the doxycycline- treated control. The mean is shown by each bar plot. N=2 biological replicates.

**Table S1:** AAV genome assemblies.

| Vector | Length | Reads | Percent of total |
| --- | --- | --- | --- |
|  | 4788 | 148 | 17.5980975 |
|  | 3866 | 122 | 14.50653983 |
|  | 2514 | 112 | 13.31747919 |
|  | 6383 | 69 | 8.20451843 |
|  | 2116 | 130 | 15.45778835 |
|  | 1071 | 109 | 12.960761 |
|  | 695 | 113 | 13.43638526 |
| 2xSURF(mU1a) | 3425 | 38 | 4.51843044 |
|  | 4837 | 150 | 32.32758621 |
|  | 2490 | 98 | 21.12068966 |
|  | 1854 | 133 | 28.6637931 |
| TwinFin | 3459 | 83 | 17.88793103 |

## References

1. van Rheenen, W., et al., Common and rare variant association analyses in amyotrophic lateral sclerosis identify 15 risk loci with distinct genetic architectures and neuron-specific biology. Nat Genet, 2021. 53(12): p. 1636–1648.

2. Dickson, D.W., K.A. Josephs, and C. Amador-Ortiz, TDP-43 in differential diagnosis of motor neuron disorders. Acta Neuropathol, 2007. 114(1): p. 71–9.

3. Arai, T., et al., TDP-43 is a component of ubiquitin-positive tau-negative inclusions in frontotemporal lobar degeneration and amyotrophic lateral sclerosis. Biochem Biophys Res Commun, 2006. 351(3): p. 602–11.

4. Neumann, M., et al., Ubiquitinated TDP-43 in Frontotemporal Lobar Degeneration and Amyotrophic Lateral Sclerosis. Science, 2006. 314(5796): p. 130–133.

5. Amador-Ortiz, C., et al., TDP-43 immunoreactivity in hippocampal sclerosis and Alzheimer’s disease. Ann Neurol, 2007. 61(5): p. 435–45.

6. Arai, T., et al., Phosphorylated TDP-43 in Alzheimer’s disease and dementia with Lewy bodies. Acta Neuropathol, 2009. 117(2): p. 125–36.

7. Josephs, K.A., et al., Abnormal TDP-43 immunoreactivity in AD modifies clinicopathologic and radiologic phenotype. Neurology, 2008. 70(19 Pt 2): p. 1850–7.

8. Kadokura, A., et al., Regional distribution of TDP-43 inclusions in Alzheimer disease (AD) brains: their relation to AD common pathology. Neuropathology, 2009. 29(5): p. 566–73.

9. King, A., et al., Abnormal TDP-43 expression is identified in the neocortex in cases of dementia pugilistica, but is mainly confined to the limbic system when identified in high and moderate stages of Alzheimer’s disease. Neuropathology, 2010. 30(4): p. 408–19.

10. Lippa, C.F., et al., Transactive response DNA-binding protein 43 burden in familial Alzheimer disease and Down syndrome. Arch Neurol, 2009. 66(12): p. 1483–8.

11. Uryu, K., et al., Concomitant TAR-DNA-binding protein 43 pathology is present in Alzheimer disease and corticobasal degeneration but not in other tauopathies. J Neuropathol Exp Neurol, 2008. 67(6): p. 555–64.

12. Agra Almeida Quadros, A.R., et al., Cryptic splicing of stathmin-2 and UNC13A mRNAs is a pathological hallmark of TDP-43-associated Alzheimer’s disease. Acta Neuropathologica, 2024. 147(1): p. 9.

13. Yamashita, R., et al., TDP-43 Proteinopathy Presenting with Typical Symptoms of Parkinson’s Disease. Mov Disord, 2022. 37(7): p. 1561–1563.

14. Tiloca, C., et al., TARDBP mutations in a cohort of Italian patients with Parkinson’s disease and atypical parkinsonisms. Front Aging Neurosci, 2022. 14: p. 1020948.

15. Halliday, G., et al., Mechanisms of disease in frontotemporal lobar degeneration: gain of function versus loss of function effects. Acta Neuropathol, 2012. 124(3): p. 373–82.

16. Ling, S.-C., M. Polymenidou, and Don, Converging Mechanisms in ALS and FTD: Disrupted RNA and Protein Homeostasis. Neuron, 2013. 79(3): p. 416–438.

17. Purice, M.D. and J.P. Taylor, Linking hnRNP Function to ALS and FTD Pathology. Front Neurosci, 2018. 12: p. 326.

18. Meneses, A., et al., TDP-43 Pathology in Alzheimer’s Disease. Mol Neurodegener, 2021. 16(1): p. 84.

19. Chung, M., et al., Cryptic exon inclusion is a molecular signature of LATE-NC in aging brains. Acta Neuropathol, 2024. 147(1): p. 29.

20. Decker, L., S. Menge, and A. Freischmidt, Cryptic exon inclusion in TDP-43 proteinopathies: opportunities and challenges. Neural Regen Res, 2025. 20(7): p. 2003–2004.

21. Ling, J.P., et al., TDP-43 repression of nonconserved cryptic exons is compromised in ALS-FTD. Science, 2015. 349(6248): p. 650–5.

22. Jeong, Y.H., et al., Tdp-43 cryptic exons are highly variable between cell types. Mol Neurodegener, 2017. 12(1): p. 13.

23. Sun, M., et al., Cryptic exon incorporation occurs in Alzheimer’s brain lacking TDP-43 inclusion but exhibiting nuclear clearance of TDP-43. Acta Neuropathol, 2017. 133(6): p. 923–931.

24. Brown, A.-L., et al., TDP-43 loss and ALS-risk SNPs drive mis-splicing and depletion of UNC13A. Nature, 2022. 603(7899): p. 131–137.

25. Ma, X.R., et al., TDP-43 represses cryptic exon inclusion in the FTD–ALS gene UNC13A. Nature, 2022. 603(7899): p. 124–130.

26. Seddighi, S., et al., Mis-spliced transcripts generate de novo proteins in TDP-43–related ALS/FTD. Science Translational Medicine, 2024. 16(734): p. eadg7162.

27. Irwin, K.E., et al., A fluid biomarker reveals loss of TDP-43 splicing repression in presymptomatic ALS–FTD. Nature Medicine, 2024. 30(2): p. 382–393.

28. Melamed, Z., et al., Premature polyadenylation-mediated loss of stathmin-2 is a hallmark of TDP-43-dependent neurodegeneration Nat. Neurosci., 2019. 22: p. 180–190.

29. Klim, J.R., et al., ALS-implicated protein TDP-43 sustains levels of STMN2, a mediator of motor neuron growth and repair. Nat Neurosci, 2019. 22(2): p. 167–179.

30. Lopez-Erauskin, J., et al., Stathmin-2 loss leads to neurofilament-dependent axonal collapse driving motor and sensory denervation. Nat Neurosci, 2024. 27(1): p. 34–47.

31. Beccari, M.S., et al., Stathmin-2 enhances motor axon regeneration after injury independent of its binding to tubulin. Proceedings of the National Academy of Sciences, 2025. 122(21): p. e2502294122.

32. Guerra San Juan, I., et al., Loss of mouse Stmn2 function causes motor neuropathy. Neuron, 2022. 110(10): p. 1671–1688.e6.

33. Baughn, M.W., et al., Mechanism of STMN2 cryptic splice-polyadenylation and its correction for TDP-43 proteinopathies. Science, 2023. 379(6637): p. 1140–1149.

34. van Es, M.A., et al., Genome-wide association study identifies 19p13.3 (UNC13A) and 9p21.2 as susceptibility loci for sporadic amyotrophic lateral sclerosis. Nat Genet, 2009. 41(10): p. 1083–7.

35. Diekstra, F.P., et al., C9orf72 and UNC13A are shared risk loci for amyotrophic lateral sclerosis and frontotemporal dementia: a genome-wide meta-analysis. Ann Neurol, 2014. 76(1): p. 120–33.

36. Finkel, R.S., et al., Treatment of infantile-onset spinal muscular atrophy with nusinersen: a phase 2, open-label, dose-escalation study. Lancet, 2016. 388(10063): p. 3017–3026.

37. Hua, Y., et al., Antisense masking of an hnRNP A1/A2 intronic splicing silencer corrects SMN2 splicing in transgenic mice. Am J Hum Genet, 2008. 82(4): p. 834–48.

38. Bennett, C.F., A.R. Krainer, and D.W. Cleveland, Antisense Oligonucleotide Therapies for Neurodegenerative Diseases. Annu Rev Neurosci, 2019. 42: p. 385–406.

39. Keuss, M.J., et al., Loss of TDP-43 induces synaptic dysfunction that is rescued by UNC13A splice-switching ASOs. bioRxiv, 2024.

40. Lesman, D., et al., U7 snRNA, a Small RNA with a Big Impact in Gene Therapy. Hum Gene Ther, 2021. 32(21-22): p. 1317–1329.

41. Gorman, L., et al., Stable alteration of pre-mRNA splicing patterns by modified U7 small nuclear RNAs. Proc Natl Acad Sci U S A, 1998. 95(9): p. 4929–34.

42. Gadgil, A. and K.D. Raczynska, U7 snRNA: A tool for gene therapy. J Gene Med, 2021. 23(4): p. e3321.

43. d’Arqom, A., et al., Engineered U7 Small Nuclear RNA Restores Correct β-Globin Pre-mRNA Splicing in Mouse β(IVS2-654)-Thalassemic Erythroid Progenitor Cells. Hum Gene Ther, 2021. 32(9-10): p. 473–480.

44. Goyenvalle, A., et al., Engineering multiple U7snRNA constructs to induce single and multiexon-skipping for Duchenne muscular dystrophy. Mol Ther, 2012. 20(6): p. 1212–21.

45. Odermatt, P., et al., Somatic Therapy of a Mouse SMA Model with a U7 snRNA Gene Correcting SMN2 Splicing. Mol Ther, 2016. 24(10): p. 1797–1805.

46. Smargon, A.A., et al., Small nuclear RNAs enhance protein-free RNA-programmable base conversion on mammalian coding transcripts. bioRxiv, 2024.

47. Byrne, S.M., et al., An engineered U7 small nuclear RNA scaffold greatly increases ADAR-mediated programmable RNA base editing. Nat Commun, 2025. 16(1): p. 4860.

48. McCarty, D.M., Self-complementary AAV vectors; advances and applications. Mol Ther, 2008. 16(10): p. 1648–56.

49. Waldrop, M. AAV9 U7snRNA Gene Therapy to Treat Boys With DMD Exon 2 Duplications. [Clinical Trial] 2023 February 9, 2023 [cited 2025 August 18]; Available from: https://www.clinicaltrials.gov/study/NCT04240314?a=5.

50. Dystrophy, P.P.M. Nationwide Children’s Announces Restoration of Full-Length Dystrophin Using Dup 2 Gene Therapy Approach. [Article] 2022 May 17, 2022 [cited 2025 August 18].

51. Hatch, S.T., A.A. Smargon, and G.W. Yeo, Engineered U1 snRNAs to modulate alternatively spliced exons. Methods, 2022. 205: p. 140–148.

52. Gorman, L., D.R. Mercatante, and R. Kole, Restoration of correct splicing of thalassemic beta-globin pre-mRNA by modified U1 snRNAs. J Biol Chem, 2000. 275(46): p. 35914–9.

53. Odermatt, P., et al., Somatic Therapy of a Mouse SMA Model with a U7 snRNA Gene Correcting SMN2 Splicing. Mol Ther, 2016. 24(10): p. 1797–1805.

54. Jia, Y., J.C. Mu, and S.L. Ackerman, Mutation of a U2 snRNA gene causes global disruption of alternative splicing and neurodegeneration. Cell, 2012. 148(1-2): p. 296–308.

55. Gunderson, S.I., M.W. Knuth, and R.R. Burgess, The human U1 snRNA promoter correctly initiates transcription in vitro and is activated by PSE1. Genes Dev, 1990. 4(12a): p. 2048–60.

56. Mangin, M., M. Ares, Jr., and A.M. Weiner, Human U2 small nuclear RNA genes contain an upstream enhancer. EMBO J, 1986. 5(5): p. 987–95.

57. Wang, Y., et al., rMATS-turbo: an efficient and flexible computational tool for alternative splicing analysis of large-scale RNA-seq data. Nat Protoc, 2024. 19(4): p. 1083–1104.

58. Vaquero-Garcia, J., et al., RNA splicing analysis using heterogeneous and large RNA-seq datasets. Nat Commun, 2023. 14(1): p. 1230.

59. Domenger, C., et al., RNA-Seq Analysis of an Antisense Sequence Optimized for Exon Skipping in Duchenne Patients Reveals No Off-Target Effect. Mol Ther Nucleic Acids, 2018. 10: p. 277–291.

60. Gushchina, L.V., et al., Lack of Toxicity in Nonhuman Primates Receiving Clinically Relevant Doses of an AAV9.U7snRNA Vector Designed to Induce DMD Exon 2 Skipping. Hum Gene Ther, 2021. 32(17-18): p. 882–894.

61. Armas, J.M.B., et al., Challenges of modelling TDP-43 pathology in mice. Mamm Genome, 2025. 36(2): p. 465–481.

62. Torregrosa, T., et al., Use of CRISPR/Cas9-mediated disruption of CNS cell type genes to profile transduction of AAV by neonatal intracerebroventricular delivery in mice. Gene Ther, 2021. 28(7-8): p. 456–468.

63. Kosaka, M., et al., Evaluation of the loading capacity and patterns of packaged DNA in AAV genomes of different sizes using long-read sequencing. Mol Ther Methods Clin Dev, 2025. 33(2): p. 101474.

64. Ayuso, E., Manufacturing of recombinant adeno-associated viral vectors: new technologies are welcome. Mol Ther Methods Clin Dev, 2016. 3: p. 15049.

65. Tran, N.T., et al., AAV-Genome Population Sequencing of Vectors Packaging CRISPR Components Reveals Design-Influenced Heterogeneity. Mol Ther Methods Clin Dev, 2020. 18: p. 639–651.

66. Uchikawa, H., et al., U7 snRNA-mediated correction of aberrant splicing caused by activation of cryptic splice sites. J Hum Genet, 2007. 52(11): p. 891–897.

67. Wang, J.H., et al., Adeno-associated virus as a delivery vector for gene therapy of human diseases. Signal Transduct Target Ther, 2024. 9(1): p. 78.

68. van der Walt, S., et al., scikit-image: image processing in Python. PeerJ, 2014. 2: p. e453.

69. Stringer, C., et al., Cellpose: a generalist algorithm for cellular segmentation. Nat Methods, 2021. 18(1): p. 100–106.

